# A pH-dependent protein kinase cascade regulates divergent differentiation of *Leishmania* in the sand fly

**DOI:** 10.64898/2026.09.15.751656

**Authors:** Laryssa V. de Liz, Jovana Sádlová, Barbora Bečvářová, James A. Brannigan, Sarah Forrester, Charlotte McNiven, Adam Dowle, Chris Taylor, Anthony J. Wilkinson, Petr Volf, Jeremy C. Mottram, Nicola Baker

**Author notes:** Corresponding author: Nicola Baker.

## Abstract

*Leishmania* parasites must rapidly adapt to fluctuating environments to ensure survival and transmission. While acidic pH in the sand fly vector is a conserved developmental trigger, sensing mechanisms remain poorly understood. Using a barcoded protein kinase library, we screened for regulators of acid adaptation in *Leishmania mexicana*, identifying nine protein kinases influencing survival at low pH, including a haptomonad differentiation regulator protein kinase (HDRK1). We demonstrate that HDRK1 null mutants *(Δhdrk1)* are predisposed to differentiate to haptomonad-like forms at low pH. While *Δhdrk1* mutants successfully infect the sand fly midgut, they fail to colonise the stomodeal valve, compromising transmission. Integrated transcriptomic and proteomic analyses revealed that at low pH, *Δhdrk1* mutants enter a low energy state reminiscent of AMPK-activated cells. We identified a second protein kinase, HDRK2, establishing a pH-dependent signalling pathway that governs the developmental fate of the parasite, directing differentiation towards either mammalian-infective metacyclic or vector-attached haptomonad stages. Finally, our screen revealed that phosphoinositide balance, regulated by the lipid kinases PI4K and PI4P5K, is vital for acid adaptation, and identified two STE transmembrane kinases as potential pH sensors. Together these findings provide a framework for how *Leishmania* detects and survives acid stress to coordinate its life cycle.

## Introduction

All organisms must adapt to environmental changes to ensure appropriate physiological or behavioural responses. Vector-borne parasites, like *Leishmania*, are an extreme example, encountering major changes in pH, temperature and nutrients upon transmission between insect and mammalian hosts. Continuous sensing and adaptation allow *Leishmania* transmission from sand flies to humans, causing the leishmaniases.

Upon sand fly ingestion, *Leishmania* amastigotes shift from 37°C to ambient temperature (26°C), and from mammalian macrophage acidity (pH 5.5) to the alkaline insect midgut (pH 8.2) (Fig.1a). These changes trigger differentiation into dividing procyclic promastigotes.^1^ During blood meal digestion, the midgut pH decreases to pH 7, prompting differentiation into an elongated nectomonad form. These parasites escape through the degraded peritrophic matrix and attach to the midgut epithelium to avoid elimination during defecation. After digestion, cells migrate to the acidic thoracic midgut (pH 5.5-6).^1,2^ Here, *Leishmania* differentiates into one of two forms. By day 8 post-infection, metacyclic promastigotes account for around 40-60% of the population and are specialised for human infection.^3,4^ Uninoculated metacyclic forms can back-transform into leptomonads, proliferate and subsequently differentiate into a new generation of metacyclic promastigotes.^5^ Conversely, haptomonad forms account for less than 10% of the population.^6^ They attach to and damage the stomodeal valve, facilitating the regurgitation and transmission of metacyclic promastigotes.^7^ Recently, the haptomonad form has also been shown to play a role in macrophage infections, although the underlying mechanism is unclear.^5^

**Figure 1.**
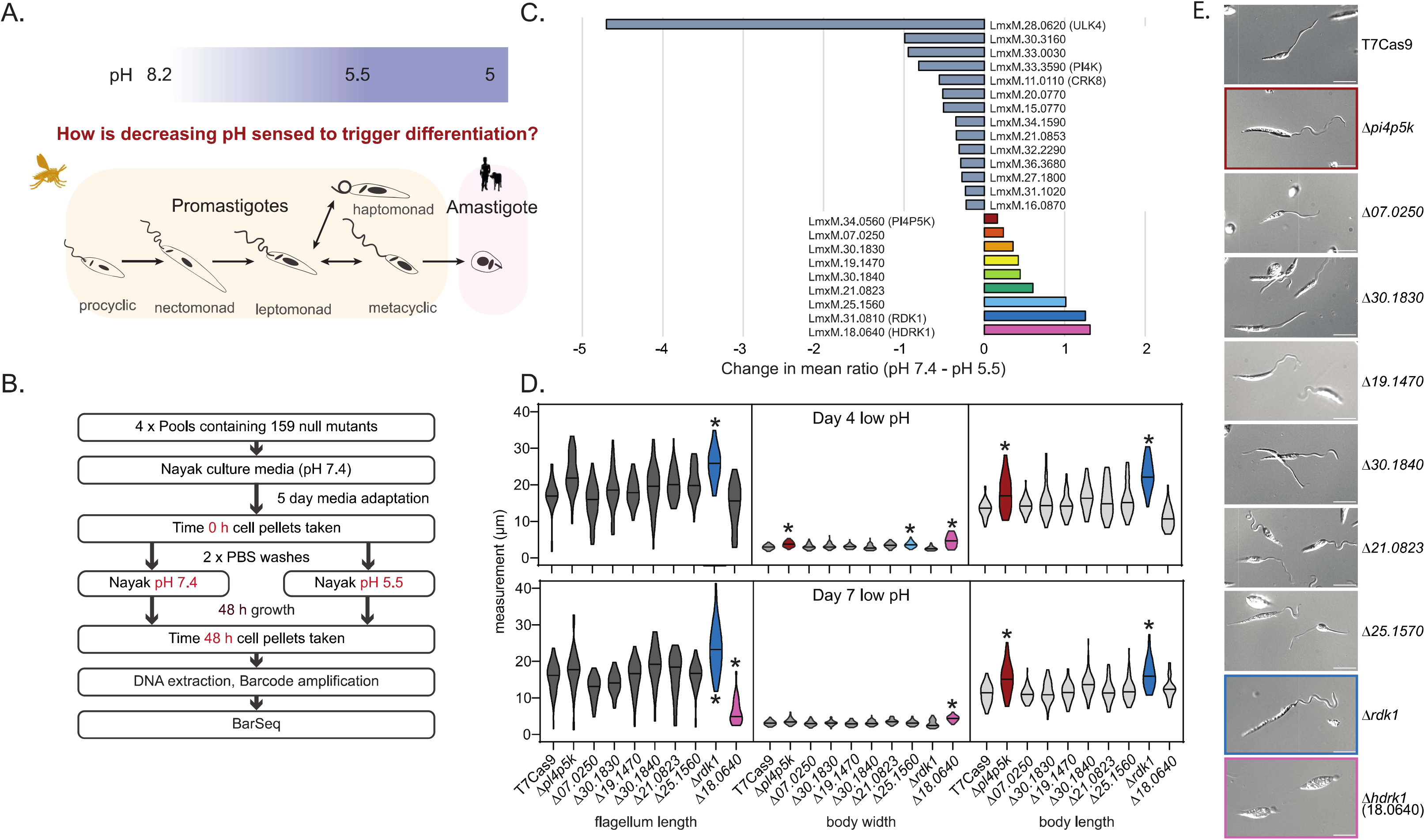
A kinome wide screen identifies protein kinases whose deletion enhances survival at low pH. A) Schematic of the *Leishmania mexicana* lifecycle stages, with the approximate pH encountered at each stage indicated. B) Schematic of the kinome-wide screen. Four replicates, each containing 159 pooled null barcoded mutants, were adapted to a Nayak medium, pH 7.4, for 5 days. Cells were washed and transferred to Nayak media at either pH 5.5 or pH 7.4 for a further 48 h. DNA was extracted from the library, and barcodes were PCR-amplified and sequenced. C) Bar chart showing mutants with a significant change in mean barcode counts between pH 7.4 and 5.5. Barcode counts from four replicates at 48 h in these two conditions were normalised against the time 0 dataset. Difference in fitness was assessed using multiple unpaired t-tests with Welch’s correction. Outputs with q-value <0.001 were plotted. Hits with a negative change in mean, indicating low fitness at pH 5.5, are shown in grey, and those with a positive change, indicating increased fitness at pH 5.5, are shown in colour. D) Phenotypic measurements, flagellum length, body width and body length, of mutant lines after being placed in Grace’s media, pH 5.5, at day 4 and day 7. Violin plots show the measurements from 50 cells, taken across two replicate experiments. Stars indicate a significant difference from the T7Cas9 parental line, P < 0.0001, following one-way ANOVA with Dunnett’s multiple comparisons test. *Δpi4p5k* is highlighted in red, *Δrdkl* is highlighted in blue and *Δhdrkl* is highlighted in magenta. E) Phase-contrast image of T7Cas9 parental cell and mutants identified with a positive difference in mean at day 7 in Grace’s media. Scale bar, 5 µm. *Δpi4p5k, Δrdkl* and *Δhdrkl* are highlighted.

Environmental pH cues drive differentiation between lifecycle stages across human-infective parasites like *Leishmania, Toxoplasma* and *Plasmodium*,^8−10^ but the signalling pathways and sensing mechanisms remain elusive. Human sensory protein families such as G-protein-coupled receptors, receptor tyrosine kinases and heat-shock transcription factors lack orthologues in *Leishmania*. This does not mean the parasite lacks proteins with equivalent functions;^11^ rather, these remain undiscovered in *Leishmania*,^12^ which likely relies on unique biology to sense environmental changes.

Protein kinases play a central role in intracellular signalling and differentiation. In *Leishmania*, the kinome is specialised, featuring expansions in CMGC, NEK and STE kinase groups.^13^ These expansions underpin bespoke signal transduction systems, as evidenced by the identification of protein kinases with sensory roles in related trypanosomes.^14^ For instance, members of the expanded CMGC family, such as DYRK kinases in *Trypanosoma brucei*, act as crucial integrators of environmental cues to drive life cycle differentiation.^15^ Many kinases have been linked to differentiation between life cycle stages.^16^ For example, RDK1 (Repressor of Differentiation Kinase 1) critically regulates the amastigote-to-promastigote transition,^17^ while protein kinase A (PKA) mediates the promastigote-to-amastigote transition in response to concurrent pH and temperature changes.^18^

Sensing pH change is not unique to life cycle transitions. Insect-stage trypanosomes follow pH gradients via taxis to navigate within the tsetse fly, metabolising glucose to acidify their environment and following these gradients via cAMP signalling.^19^ The specific pathways required to sense pH change remain elusive.

We used a barcoded *Leishmania* protein kinase deletion library to systematically identify protein kinases that mediate adaptation to acidic environments. Here, we characterise HDRK1, a CAMK-family kinase required for leptomonad-to-metacyclic promastigote differentiation. We demonstrate that without HDRK1, parasites fail to complete metacyclogenesis and instead divert into a haptomonad-like state. Finally, we consider how HDRK1 integrates into a wider signalling network of pH-responsive protein kinases identified in our screen.

## RESULTS

### A kinome screen identifies nine *Leishmania* protein kinases regulating low-pH survival

To identify protein kinases essential for surviving environmental transitions in the *Leishmania* life cycle, we utilised a kinome-focused barcoded knockout library of 159 mutants^16^ adapted to defined Nayak medium.^20^ Post-adaptation, pooled mutants were placed at pH 7.4 or pH 5.5 for 48 h. BarSeq analysis determined relative abundance ratios (normalised T0/T48 reads)(Fig. 1b), assigning the highest values to the most depleted mutants. Multiple pairwise t-tests tracked changes in mean ratio (Fig. 1c, Supplementary Table 1). A positive change in mean ratio identifies mutants overrepresented at pH 5.5 relative to pH 7.4 (negative regulators of low-pH survival), whereas a negative change identifies mutants required for low-pH survival.

Pairwise comparisons revealed 23 kinases with significant fitness alterations under low-pH. 14 kinases were depleted at pH 5.5, identifying them as required for survival at pH 5.5. These included Unc-51-like kinase 4 (ULK4), required for flagellar cytoskeleton assembly^21^, phosphatidylinositol 4-kinase (PI4K), and members of the NEK (LmxM.21.0853, LmxM.30.3160 and LmxM.34.5190), STE (LmxM.32.2290, LmxM.36.3680, LmxM.31.1020 and LmxM.20.0770), NAK (LmxM.15.0770 and LmxM.33.0030), CMGC (LmxM.11.0110 and LmxM.27.1800) and orphan (LmxM.16.0870) kinase families. PI4K and ULK4 were previously implicated in sand fly midgut colonisation^16^, linking acid tolerance to survival in the vector. Conversely, 9 null mutants gained representation at pH 5.5, identifying these kinases as negative regulators of low-pH adaptation.

We investigated the nine null mutants with a survival advantage at pH 5.5 to uncover molecular mechanisms restricting acidic adaptation rather than general survival. As incubation in acidic medium for 7 days induces differentiation into metacyclic forms^22^, we performed morphological profiling (flagellar length, cell body width/length) of these mutants in pH 5.5 Grace’s insect medium. This identified a striking phenotype for the LmxM.18.0640 mutant (Fig. 1d). By day 7, rather than transforming into metacyclic forms, over 85% of the population developed an “extra-axonemal bulb”, characteristic of haptomonads (Fig. 1e). We named this gene Haptomonad Differentiation Regulator Kinase 1 (HDRK1).

Morphological profiling also identified body length increases in *Δrdkl* and *Δpi4p5k*, at days 4 and 7 in Grace’s medium. *Ardkl* additionally showed significantly longer flagellar relative to parental T7Cas9. Life cycle stage profiling^2^ showed that 72% of *Δrdk1* and 58% of *Δpi4p5k* cells displayed elongated nectomonad morphology, compared to 14% in T7Cas9. In contrast, metacyclic forms were absent in *Δhdrk1*, compared to 6% seen in T7Cas9, highlighting its divergence from normal metacyclogenesis (Supplementary information).

### HDRK1 activity mediates differentiation into the metacyclic promastigote stage

We imaged *Δhdrk1* cells grown in Grace’s media daily to track extra-axonemal material (flagellar membrane swelling). After 24 h at low pH, almost half the population displayed this phenotype, exceeding 90% by day 7. The material starts as a small swelling that becomes a bulb, accompanied by flagellar shortening, reminiscent of a haptomonad-like phenotype (Fig. 2a). We hypothesised that HDRK1 disruption prevents metacyclic promastigote differentiation, committing cells to a haptomonad-like fate.

**Figure 2.**
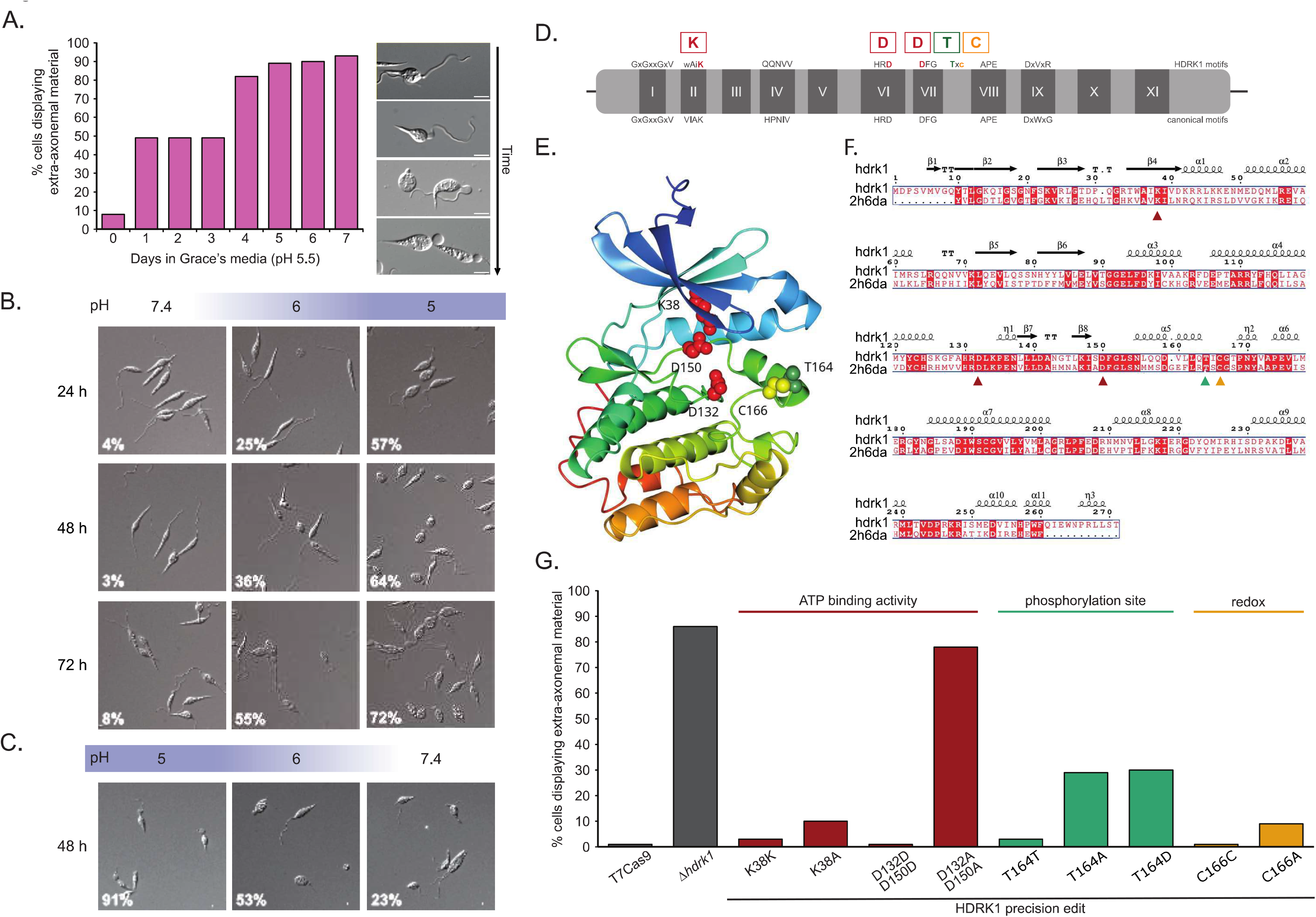
HDRK1 is an active kinase required for differentiation into the metacyclic promastigotes. A) Bar chart indicating the percentage of population with extra-axonemal material over 7 days in Grace’s media (n >86). Day 0 represents cells before transfer to pH 5.5. Phase-contrast images show examples of cells with the extra-axonemal material. Scale bars, 5 µm. B) Phase-contrast images of mid-log*Δhdrkl* procyclic promastigotes washed and grown in HOMEM medium at differing pH values, pH 7.4, 6.0 and 5.0, for 24, 48 and 72 h. The percentage of cells with extra-axonemal material indicated in white text. C) Phase-contrast images of *Δhdrkl* cells grown for 7 days at pH 5.5, then washed and resuspended in HOMEM medium at pH 5.0, 6.0 or 7.4 for 48 h. Scale bars, 5 µm. The percentage of cells with extra-axonemal material is indicated in white text. D) Schematic of the HDRK1 (LmxM.18.0640) protein kinase, with the conserved motifs depicted below. Residues of interest are indicated: residues required for kinase catalytic activity, lysine and aspartic acid, are shown in red; T164, a confirmed phosphosite, is shown in green; and C166, a cysteine, linked to redox activation, is shown in yellow. E) Ribbon diagram of the HDRK1 AlphaFold model with residues of interest highlighted. K38, D132 and D150 are shown in red, T164 in green and C166 in yellow. F) Sequence alignment of HDRK1 and the kinase domain of human AMPKα (AAPK2) in the context of secondary structure elements from the HDRK1 AlphaFold model. G) Bar chart showing the percentage of population with extra-axonemal material for T7Cas9 and HDRK1 precision-edited lines, K38, D132/D150, T164 and C166, grown at low pH for 7 days.

To characterise HDRK1, clonal null mutants with a single *BLA* antibiotic marker, *Δhdrk1*, were generated together with an add-back line expressing HDRK1 from a ribosomal locus, *△hdrk1*/HDRK1(Supplementary Fig. 1). Mutants were grown in HOMEM medium (pH 7.4, pH 6.0 or pH 5.0) to test whether acidity alone induces bulb formation (Fig. 2b). At pH 7.4, low levels of cells exhibited extra-axonemal material (4% at 24 h), but acidic medium caused the phenotype to increase over time and with lower pH. At pH 6, extra-axonemal material developed in 25% of the population after 24 h, progressing to 36% after 48 h and 55% by 72 h. This was more pronounced at pH 5, with 57%, 64% and 72% of the population exhibiting extra-axonemal material at 24, 48 and 72 h, respectively. When cells grown at pH 5.5 for 7 days were transferred to medium at pH 5, pH 6 or pH 7.4 for 2 days, the phenotype was reversible, with 91% of the population at pH 5 retaining extra-axonemal material compared to 53% at pH 6 and 23% at pH 7.4 (Fig. 2c). Flagellar remodelling appears to start before 48 h (Supplementary Fig. 2a). *△hdrk1*/HDRK1 cells behave like T7Cas9, confirming that ribosomal locus expression complements the mutation (Supplementary Fig. 1c).

### HDRK1 is an AMPK/SNF1-like kinase dependent on catalytic activity

HDRK1 belongs to the Ca^2+^ /calmodulin-dependent protein kinase (CAMK) family, which includes AMP-activated protein kinase (AMPK) and yeast sucrose non-fermenting (SNF1) orthologues linked by a shared upstream CAMKK protein.^23^ SNF1 and AMPK are important metabolic sensors responsible for maintaining cellular energy balance.^24^ HDRK1 comprises a kinase domain without a defined calmodulin domain (Fig. 2d). Overlaying an HDRK1 AlphaFold model (Fig. 2e) with the human AMPKα kinase crystal structure reveals high structural similarity (rmsA - 1.6 Å for 248 matched Ca atoms) and sequence conservation (Fig. 2f). Structural differences span residues 152-169, containing the activation loop, which is expected to be conformationally flexible and therefore show low AlphaFold confidence.

SNF1 and AMPKα are activated via threonine phosphorylation in the activation loop,^25^ conserved as T164 in HDRK1 (Fig. 2f).^26^ HDRK1 has three catalytic residues (K38, D132, D150) and a cysteine (C166) in a T+2 position which facilitates redox-mediated activation in other kinases with a cysteine at this position (Fig. 2d, 2e, 2f).^13,27^ HDRK1 expressed and purified from bacteria exhibited >5-fold increased *in vitro* activity against a peptide containing the conserved AMPK/SNF1 substrate motif (CD-x-R-X-X-S-x-x-x-Φ; Φ = hydrophobic residue) using an ADP-Glo assay (Supplementary Fig. 2b,c).^28^

To determine whether catalytic activity is required for HDRK1 function, we used CRISPR-Cas9 precision editing to create mutants with compromised kinase activity.^29,30^ Alanine substitutions targeted lysine 38 and aspartates 132 and 150, which normally stabilise MgATP or promote phosphotransfer (Supplementary Fig. 3). After 7 days at low pH, the silent control mutants HDRK1^K38K^ and HDRK1^D132D,D150D^ phenocopied T7Cas9 cells, while the HDRK1^K38A^ mutant showed a mild phenotype (10% of cells having the bulb structure). This mild phenotype, paired with the unusual VIAK/HPNV motifs (WAIK/QQNVV), suggests that HDRK1 uses an atypical mode of ATP stabilisation. In contrast, the HDRK1^D132A/D150A^ double mutant phenocopied *Δhdrk1*, confirming that HDRK1 kinase activity is required to prevent haptomonad differentiation and permit metacyclogenesis (Fig. 2g, Supplementary Fig. 2d).

We next targeted the potential T164 phosphosite and putative redox-sensitive C166 cysteine. While the silent controls phenocopied T7Cas9 cells, the non-phosphorylatable HDRK1^T164A^ and potentially phosphomimetic HDRK1^T164D^ mutants both exhibited ~30% of cells with bulb structures (Fig. 2g, Supplementary Fig. 2d). These data suggest phosphorylation of T164 is important for regulation of HDRK1 function, but other regulatory mechanisms also exist.

### *Δhdrk1* mutants fail to attach to the sand fly stomodeal valve

T7Cas9, *Δhdrk1* and *Δhdrk1*/HDRK1 lines developed heavy infections in *Lutzomyia longipalpis* females (Fig. 3a). At day 1 post blood meal (PBM), all lines were enclosed within the endoperitrophic space, inside the peritrophic matrix. Differences emerged in late-stage infections post-defecation of the peritrophic matrix, when parasites migrated from the abdominal to the thoracic midgut. On days 4−5 PBM, T7Cas9 and ▵hdrk1/HDRK1 cells reached the stomodeal valve in most females and attached to it. Conversely, *Δhdrk1* parasites reached the cardia, the anterior midgut near the stomodeal valve, but failed to attach. This pattern intensified by day 9 PBM, when T7Cas9 and *Δ*hdrk1/HDRK1 colonised the stomodeal valve in all infected sand flies, while *Δhdrk1* filled the midgut and cardia region, without colonisation of the stomodeal valve (Fig. 3b).

**Figure 3.**
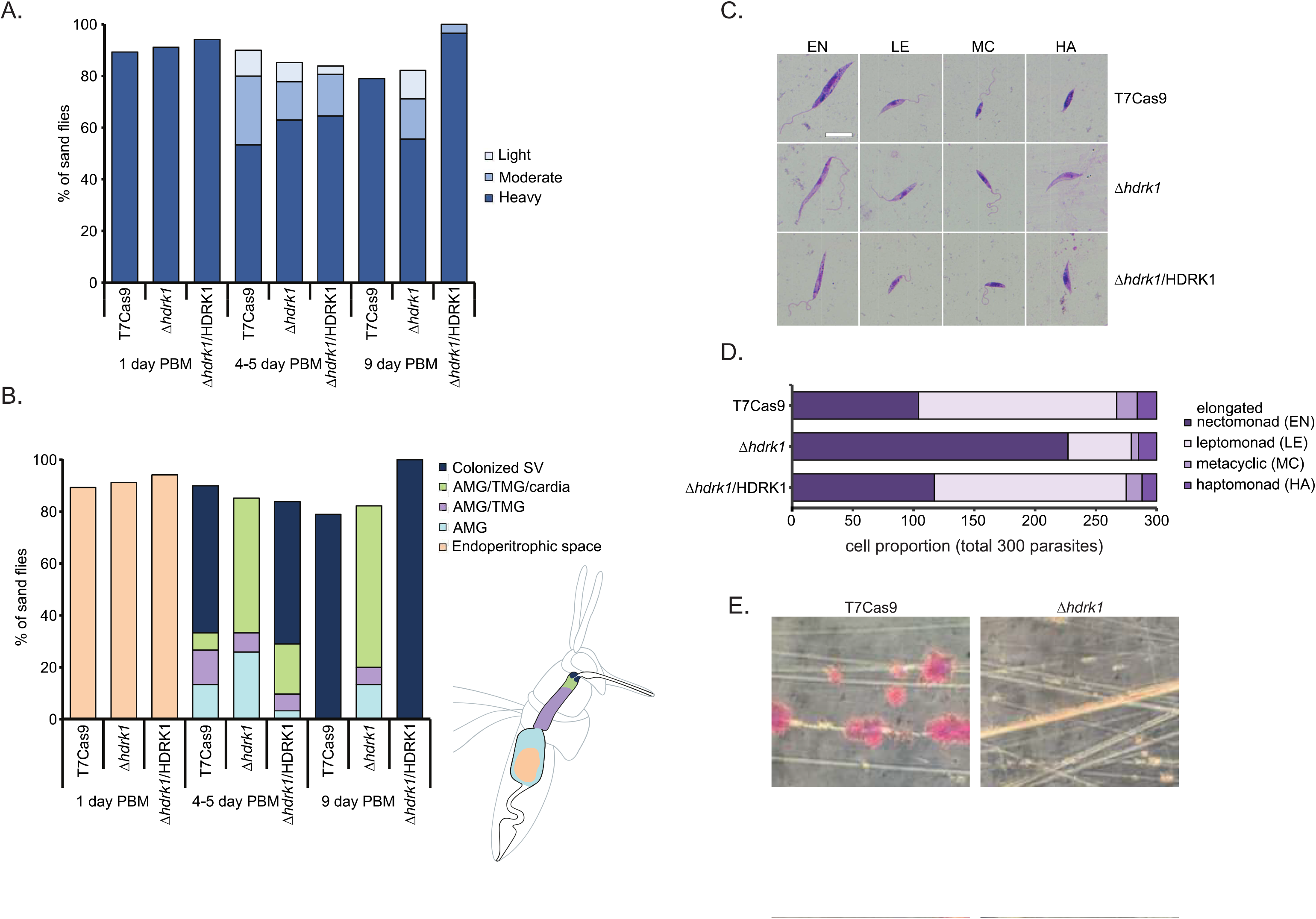
*Δhdrkl* cells cannot attach *in vitro* or *in vivo*. A) Bar chart showing infection rates and parasite loads for T7Cas9, *Δhdrkl* and *Δhdrk1*/HDRK1 in *L. longipalpis* at days 1, 4−5 and 9 post blood meal (PBM). Parasite loads were classified as light, <100 parasites per gut; moderate, 100−1,000 parasites per gut; or heavy, >1000 parasites per gut. Data were collected from two experiments, with approximately 10−20 sand fly dissections for each line at each time point. B) Bar chart showing the location of parasites in sand flies at days 1, 4−5 and 9 PBM. The sand fly schematic shows the digestive tract, colour-coded to indicate parasite location: endoperitrophic space (pink); abdominal midgut (AMG; light blue); abdominal and thoracic midgut (AMG/TMG; lilac); abdominal midgut, thoracic midgut and cardia (AMG/TMG/cardia; lime); and colonised stomodeal valve (SV; dark blue). C) Giemsa-stained cells representing the morphological forms found in the day 9 sand fly dissections for T7Cas9, *Δhdrkl* and *Δhdrk1*/HDRK1. Scale bar, 10 pm. D) Bar chart showing the relative representation of individual morphological categories in T7Cas9, *Δhdrkl* and *Δhdrk1*/HDRK1 detected in *L. longipalpis* at day 9 PBM. E) Image of scratched plastic after incubation with T7Cas9 and *Δhdrkl* mid-log phase parasites for 24 h, followed by Giemsa staining. Brightfield image acquired at 40x magnification.

*Leishmania* cells dissected from midguts on day 9 PBM had developed into all expected morphological categories in all three cell lines (elongated nectomonads, leptomonads, haptomonads and metacyclic promastigotes) (Fig. 3c). The representation of elongated nectomonads was higher in Δ*hdrk1*, whilst leptomonad and metacyclic promastigote representation was lower compared to T7Cas9 and Δhdrk1/HDRK1 (Fig. 3d). Haptomonad frequencies were comparable across lines, but *Δhdrk1* haptomonads were elongated forms with a significantly longer body and flagellum than those of T7Cas9 and âhdrk1/HDRK1 (Supplementary Fig. 4a). Elongated *Δhdrk1* haptomonads also had significantly longer bodies than leptomonads (Supplementary Fig. 4b). According to Walters^31^, these elongated haptomonads represent transitional forms of leptomonads which attach to the stomodeal valve before transforming to shorter haptomonad forms. Elongated haptomonads that developed in *Δhdrk1* were unable to attach to or colonise the stomodeal valve. These data suggest that, while *Δhdrk1* has haptomonad-like morphology, these cells are functionally defective in surface attachment, stalling the life cycle at the stomodeal valve. This attachment defect was confirmed *in vitro, as Δhdrk1* could not attach to scratched plastic as observed with T7Cas9 (Fig. 3e).

### Low pH triggers transient HDRK1 upregulation and relocation from the basal body to the flagellum

HDRK1 tagged with mNeonGreen (mNG) at its N-terminus (mNG::HDRK1) localised primarily to the basal body in cells grown at pH 7.4, with weak cytoplasmic and flagellar signals (Fig. 4a). At low pH, basal body localisation remained but flagellar signal increased after 24 h (Fig. 4b). After 4 days, the flagellar signal decreased while signal became visible in the lysosome (Supplementary Fig. 4c). In attached cells, mNG::HDRK1 localised to the attachment plaque, flagellar tip and a distinct cell body region. In detached cells, mNG::HDRK1 localised throughout the bulb structure (Fig. 4c). HDRK1 protein abundance increased over the first 48 h at low pH and then decreased sharply after 4 days, highlighting its early role in the acid response (Fig. 4d).

**Figure 4.**
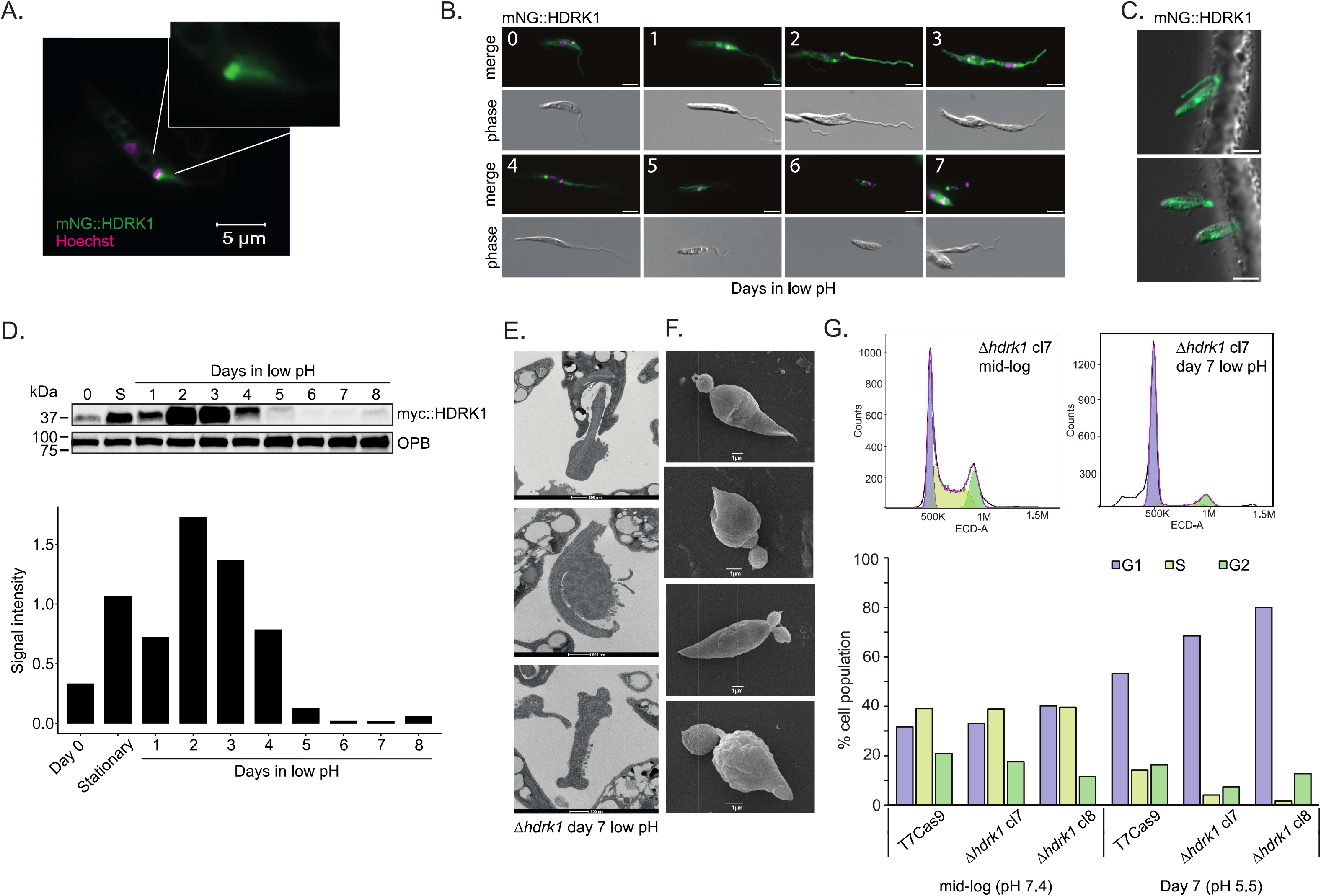
HDRK1 expression increases early in exposure to low pH. A) Live-cell image of mNG::HDRK1 in a cell from a mid-log culture grown in HOMEM medium, pH 7.4. The nucleus and kinetoplast shown in magenta. Basal body signal from mNG::HDRK1 is shown in green. Scale bar, 5 µm. B) Localisation of mNG::HDRKl in live cells grown in Grace’s media at pH 7.4 (0) or pH 5.5 for 1-7 days. Scale bar, 5 µm. C) Live-cell imaging in CyGel of mNG::HDRK1 promastigotes attached to a gridded slide. Scale bar, 5 µm. D) Expression levels of myc::HDRK1 over 8 days in low pH (Grace’s media, pH 5.5). OPB was used as a protein loading control. Day 0, mid-log cells in pH 7.4. S, stationary-phase culture in pH 7.4 media. Day 1-8, cells in low pH Grace’s media (pH 5.5). Bar chart showing HDRK1 signal intensity normalised to the OPB control. E) TEM images of *Δhdrkl* cells grown at low pH for 7 days. Scale bars, 500 nm. F) SEM images of *Δhdrkl* cells grown at low pH for 7 days. Scale bars, 1 pm. G) Flow cytometry showing the cell cycle stage (G1, S or G2) in T7Cas9 and *Δhdrkl* cells at pH 7.4 and after 7 days at pH 5.5.

Without HDRK1, extra-axonemal material accumulates in the flagellum, where mNG::HDRK1 normally translocates at low pH, and cells form a convoluted bulb structure, which may aid flagellar curling (Fig. 4e, 4f). Accordingly, multiple axoneme structures are visible in some samples as the flagellum curls and cells adopt a detached haptomonad-like morphology. Cells displaying two flagella and bulb structures could indicate that they were mid-division when the bulb structure formed, or that these haptomonad-like forms continue to divide (Fig. 4f, Supplementary Fig. 4d). Wild-type promastigotes grown at pH 5.5 stop dividing after 4 days, when metacyclic parasites are found in the culture.^22^ Flow cytometry revealed increased G1 arrest after 7 days at low pH in T7Cas9 and *Δhdrk1*, with stronger arrest in *Δhdrk1* clones, indicating that haptomonad-like forms also do not divide (Fig. 4g, Supplementary Fig. 4e).

### Transcriptomic profiling reveals signatures of haptomonad-like differentiation

We profiled transcriptomes of T7Cas9 cells and two HDRK1 clonal mutants *(Δhdrk1* cl7 and *Δhdrk1* cl8) grown for 0, 4 and 7 days in Grace’s medium (Fig. 5a, Supplementary Table 2). Principal component analysis (Fig. 5b) and hierarchical clustering (Supplementary Fig. 5a) separated T7Cas9 from *Δhdrk1* clones. Differential expression (DE) analysis compared *Δhdrk1* clones to T7Cas9 at each time point (5% FDR, log2 fold change <-1, >1) (Fig. 5c, Supplementary Fig. 5b). Across both clones, 75 upregulated and 83 downregulated transcripts were identified (Supplementary Table 3) and categorised by time point (Fig. 5d).

**Figure 5.**
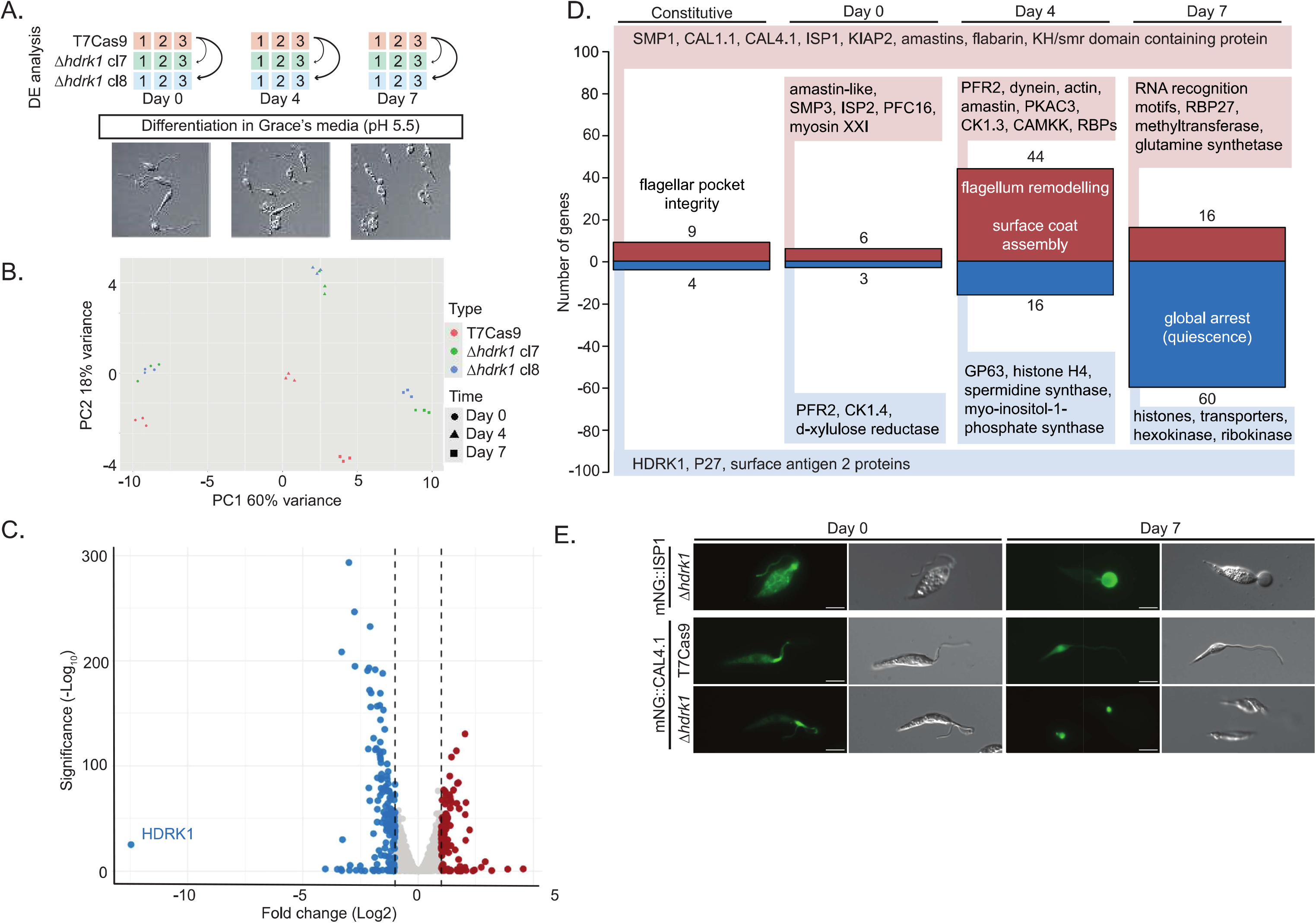
Transcriptomics reveals cellular changes in haptomonad-like forms. A) Schematic of the RNA-seq experiment. Three replicates were collected for T7Cas9, *Δhdrkl* cl7 and *Δhdrk1* cl8 cells at day 0, 4 and 7 in Grace’s media, pH 5.5. Phase-contrast images show the cell morphology of the *Δhdrkl* cl7 culture from which RNA was extracted. B) PCA plot of RNA-seq data. Triplicate datasets from T7Cas9 are shown in pink, *Δhdrkl* cl7 in green and *Δhdrkl* cl8 in blue. C) Volcano plot of differentially expressed transcripts from *Δhdrkl* cl7 compared to T7Cas9 at day 7. Transcripts with a log2 fold change <-2 are highlighted in blue, downregulated, and transcripts with a log2 fold change >2 are highlighted in red,upregulated. HDRK1 is labelled. D) Overview of transcripts differentially expressed in both *Δhdrkl* cl7 and *Δhdrkl* cl8 cells compared to T7Cas9 across the three time points (day 0, 4 and 7). Upregulated genes shown in red and downregulated genes shown in blue. E) Live-cell image of cells expressing mNG::ISP1 at day 0 (pH 7.4) and day 7 in low pH (pH 5.5). mNG::LmxM.16.0490 and mNG::Cal4.1 in T7Cas9 and *Δhdrkl* cell lines at day 0 and day 7 in Grace’s media. Scale bar, 5 µm.

Global GO analysis of the full DE transcript list showed no broad biological process enrichment but identified downregulation of nucleoside transport/salvage and three cell-adhesion proteins (Supplementary Fig. 5c). Time-resolved DE analysis revealed distinct trends (Fig. 5d). On day 4, upregulated transcripts were linked to flagellar reorganization (PFR2, dynein, tubulin, actin, PKAC3, CK1.3). Downregulated transcripts at day 4 included histone H4 and spermidine synthetase, indicative of cell-cycle arrest onset. This is more pronounced by day 7, when histones H1-4 were all downregulated, together with several transporters, hexokinase and ribokinase, suggesting that cells adopt an arrested, quiescent state.

Genes constitutively altered across all time points reveal cellular changes directly resulting from HDRK1 disruption, independent of pH change, with 9 upregulated and 4 downregulated genes (Supplementary Fig. 5d). The upregulated transcripts include three calpain-like cysteine peptidases (CALP1.1, KIAP2 and CAL4.1), two amastin surface glycoprotein genes (LmxM.16.0490 and LmxM.16.0470), flabarin (LmxM.27.1730), small myristoylated protein-1 (SMP1), inhibitor of serine peptidase 1 (ISP1, LmxM.15.0300) and a KH-domain-containing protein (LmxM.30.2810). KIAP2, SMP1 and ISP1 have previously been shown to be expressed in haptomonads.^32,33^ Furthermore, SMP1 is involved in *Leishmania* attachment within the sand fly.^34^ The downregulated genes include HDRK1, a P27 protein involved in ATP production (LmxM.28.0980) and two surface antigen genes found in a tandem array (LmxM.12.0890, LmxM.12.0891).

ISP1, a known haptomonad bulb marker,^33^ localised to extra-axonemal material (day 0) and the bulb structure (day 7) in *Δhdrk1* cells, visualised by mNG-tagging (Fig. 5e) and antibody staining (Supplementary Fig 6a). As a marker of HDRK1 disruption, we tracked CAL4.1 (LmxM.04.0450) localisation. mNG::Cal4.1 localises to the base of the flagellum in *Δhdrk1* and T7Cas9 cells at pH 7.4. After 7 days at low pH, the signal becomes weakly cytoplasmic in T7Cas9 cells and specific to the bulb structure in *Δhdrk1*. This bulb localisation is also seen in T7Cas9 cells attached to gridded glass coverslips (Supplementary Fig. 6c). Flow cytometry confirmed higher mNG::CAL4.1 expression in *Δhdrk1* versus T7Cas9 after 7 days at low pH (Supplementary Fig. 6b).

### Proteomic analysis highlights energy suppression and cytoskeletal reorganization at low pH

To complement the transcriptomic data, we compared whole-cell proteomes of *Δhdrk1* cl7 at day 0 (pH 7.4) and day 7 (pH 5.5) (Fig. 6a). Of 3833 identified proteins (5% FDR, a minimum two-peptide cutoff) (Supplementary Table 4), 441 were differentially abundant (>1 log2 fold change cutoff) (Fig. 6b, Supplementary Fig. 6d).

**Figure 6.**
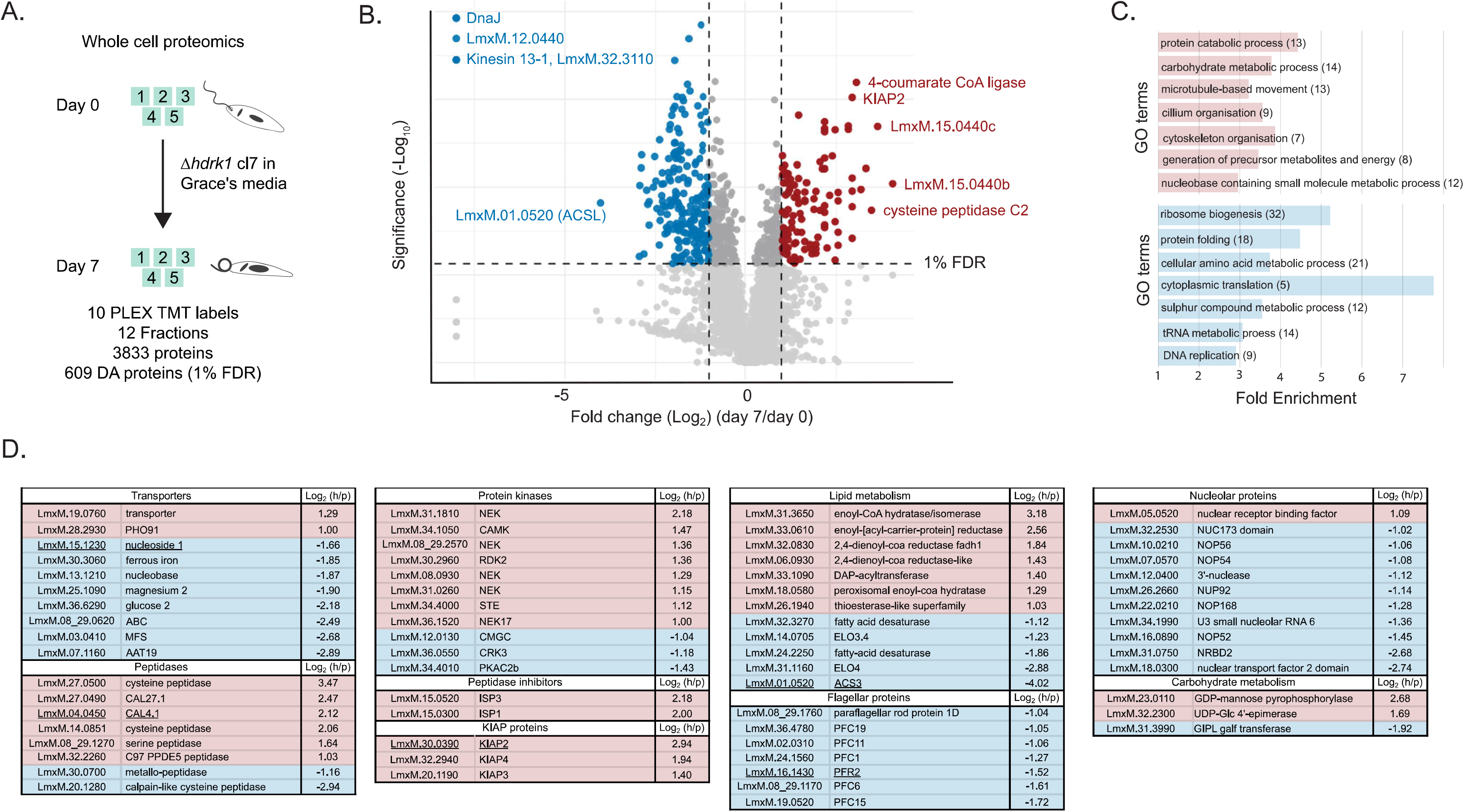
Whole-cell proteomics of haptomonad-like cells. A) Schematic of whole-cell proteomics experiment. Five replicates of *Δhdrkl* cl7 were grown in Grace’s media. Protein samples were taken at day 0 and day 7 for differential abundance analysis. B) Volcano plot showing proteins differentially abundant between day 7, pH 5.5, and day 0, pH 7.4. Blue dots represent the 275 proteins downregulated in *Δhdrkl* cl7 after 7 days at low pH, log2 fold change <-1 and <1% FDR. Red dots represent the 166 upregulated proteins after 7 days at low pH, log2 fold change >1 and <1% FDR. Selected data points with the largest log2 fold changes are labelled. C) GO term analysis results for the proteins upregulated, red, and downregulated, blue, in *Δhdrkl* haptomonad-like cells at <1% FDR. D) Table showing trends in proteins associated with shared pathways or cellular localisation. Data refers to log2 fold change of haptomonads (day 7) in comparison to promastigotes (day 0) (h/p). Red represents increased abundance, and blue represents decreased abundance. Underlined proteins were also identified as differentially expressed in the transcriptomic data.

Four of the most downregulated proteins include DnaJ (LmxM.31.3030), two hypothetical proteins (LmxM.12.0440, LmxM.32.3110) and kinesin 13-1 (LmxM.01.0030). These were not detected at day 7. The two most upregulated proteins mapped to contigs (LmxM.15.0440c and LmxM.15.0440b), linked to the accession LmxM.15.0440. Other highly upregulated proteins include the attachment protein KIAP2, cysteine peptidase 2C and 4-coumarate-CoA ligase.

GO term analysis revealed enrichment for structural remodelling and energy metabolism, specifically cytoskeleton and cilia organisation (including microtubule-based movement) and carbohydrate metabolic processes. In contrast, pathways involved in protein synthesis and assembly, as well as amino acid metabolism, were decreased (Fig. 6c, Supplementary Table 5).

Grouping proteins by cellular localisation or biological function (Fig. 6d) revealed global downregulation of flagellar proteins, nucleolar components and transporters. Conversely, peptidases, peptidase inhibitors and three KIAP proteins linked to attachment were widely upregulated in the cells at day 7 low pH. Proteins involved in fatty acid synthesis were downregulated, whilst those involved in lipid oxidation were upregulated. This was further supported by the presence of more low-molecular-weight lipids in the absence of HDRK1 on a Stains-All gel (Supplementary Fig. 7a). Carbohydrate metabolism pointed to a suppression of gluconeogenesis. Additionally, the two genes encoding malic enzyme, required for gluconeogenesis^35^, were both significantly downregulated at day 7 but below the 1-fold cutoff (Supplementary Table 4). Finally, 11 protein kinases were identified (8 upregulated, 3 downregulated). Five of the eight upregulated kinases belong to the NEK family. PKAC1, CRK3 and another CMGC family protein kinase were downregulated, perhaps due to not being required for division or motility.^36,37^

Transcriptomic trends mirrored proteomic trends. Comparison of the datasets revealed 19 genes (11 upregulated and eight downregulated) (Supplementary Fig. 7b). Notably, six of the upregulated genes were previously linked to flagellar adhesion, filament architecture or the haptomonad life cycle stage (KIAP2, SMP-1, SMP1-like, myosin XXI, ISP1, actin).^32,33,38^

### Identification and characterisation of HDRK2, a second haptomonad differentiation kinase

Null mutants of the differentially expressed kinases identified by RNA-seq or proteomics (log2 fold change >1 and <1% FDR) were retrieved,^16^ grown in Grace’s media and imaged on day 7 (Supplementary Fig. 7c). NEK17 (LmxM.36.1520) phenocopied *Δhdrk1*, with most cells forming bulb structures, and was therefore named HDRK2 (Fig. 7a). HDRK2 consists of an N-terminal kinase domain followed by a pleckstrin homology (PH) domain (Fig. 7b). The kinase domain has the three signature catalytic residues associated with ATP binding and phosphotransferase activity. PH domains are associated with proteins involved in intracellular signalling often binding to the inositol phosphate moieties of membrane phosphatidylinositol lipids.^39^

**Figure 7.**
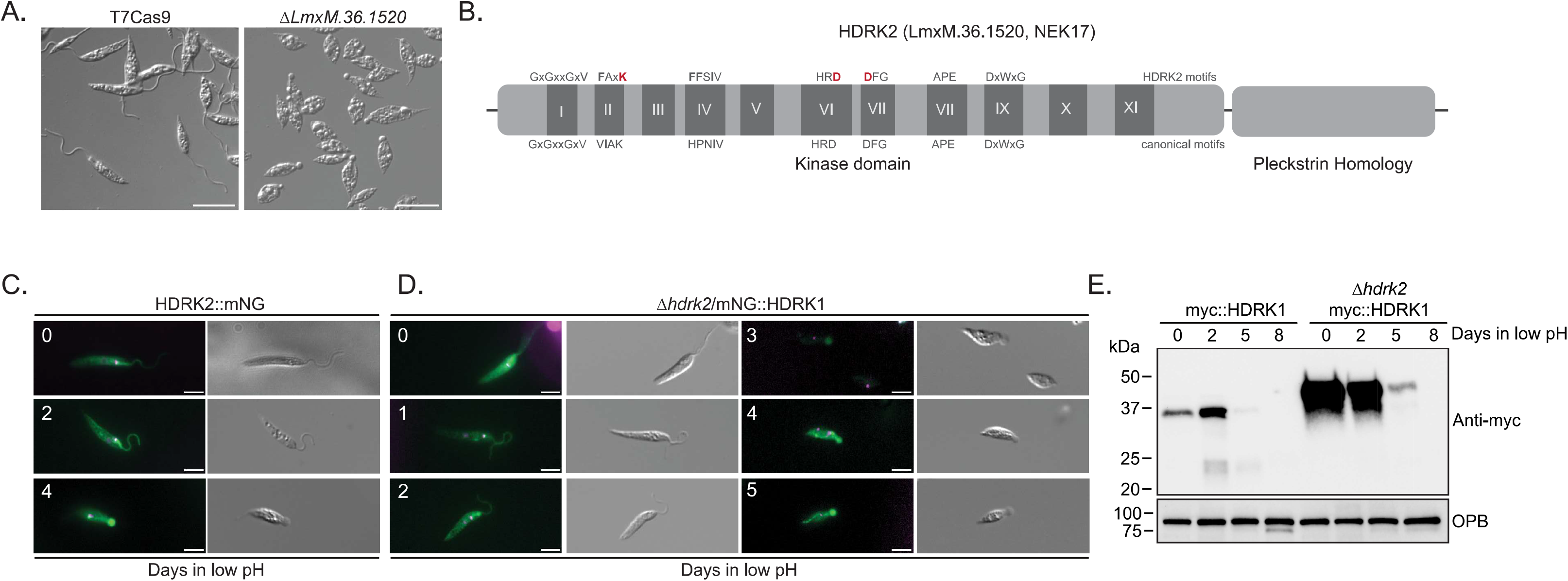
Proteomics identifies a second haptomonad differentiation kinase, HDRK2. A) Phase-contrast images of fixed T7Cas9 and *ΔLmxM.36.l520* cells after 7 days in Grace’s media. Scale bar, 10 µm. B) Schematic of HDRK2 (LmxM.36.1520). Canonical protein kinase motifs are shown below the cartoon. HDRK2-specific motifs shown above. The three residues required for kinase activity are highlighted in red. C) Live-cell images showing mNG::HDRK2, green, at days 0, 2 and 4 in low pH, Grace’s media. Hoechst is shown in magenta. D) Live-cell images of Ahdrk2/mNG::HDRK1 at days 0–5 in low pH, Grace’s media. Left, merged image with Hoechst, magenta, and mNeonGreen, green. Right, phase-contrast image. Scale bars, 5 µm. E) Western blot showing HDRK1 expression at 0 (mid-log at pH 7.4), 2, 5 and 8 days in low pH (pH 5.5) in T7Cas9 and *Δhdrk2* cells. OPB is shown as a loading control.

Weak cytoplasmic signal was detected after addition of an N-terminal mNG-tag to HDRK2 (Supplementary Fig. 8a). However, signal from the basal body region was observed after addition of a C-terminal tag (Fig. 7c, Supplementary Fig. 8a). HDRK2::mNG localised to the basal body at day 1 in low-pH medium and moved down the flagellum by day 2, as seen for HDRK1. By day 4, many haptomonad-like cells become apparent in the culture, indicating that the C-terminal tag disrupts protein function. Addition of a smaller myc epitope tag also impaired kinase activity (Supplementary Fig. 8b and 8c). Nonetheless, cells survive low pH and form a bulb structure when both HDRK1 and HDRK2 are disrupted (Supplementary Fig. 8d). HDRK1 was tagged in *Δhdrk2*, enabling the localisation on HDRK1 to be followed in a cell line that phenocopies HDRK1 deletion (Fig. 7d). The HDRK1 signal was less pronounced with higher cytoplasmic signal in *Δhdrk2*, but still moved from the basal body to the bulb structure in agreement with mNG::HDRK1 cells attached to gridded slides (Fig. 4c). Western blotting of myc::Hdrk1 in *Δhdrk2* at low pH (Fig. 7e) revealed surprisingly higher expression and stability than in T7Cas9, alongside ~8 kDa larger band, suggesting a post-translational modification.

### Functional characterisation of transmembrane kinases identified in the low-pH screen

To determine whether low-pH survival kinases share Hdrk1’s pathway, we prioritised three kinases with predicted membrane-spanning domains, characteristic of cell-surface sensors. Two tandemly arranged STE kinases, LmxM.30.1830 and LmxM.30.1840, belong to the STE family and share 57% sequence identity. This includes a central 333-residue segment with 100% sequence identity. RDK1 exhibits ~20% sequence identity with this pair. Structural analysis of models generated using AlphaFold3^40^ and domain searches in Foldseek^41^ revealed a striking similarity in the domain organisation of all three proteins. Each comprises an N-terminal dCACHE domain, a central domain with a nucleotidyl cyclase-like fold (NucCase) and a C-terminal protein kinase-like domain (Fig. 8a). The dCACHE domain is flanked by a pair of confidently predicted transmembrane sequences, consistent with extracytoplasmic localisation of this domain. dCACHE domains have widespread roles in sensing and transducing extracellular signals in bacteria, archaea and eukaryotes.^42^ Notable examples are McpA and TlpA from *Bacillus subtilis*, whose dCACHE domains sense extracellular pH as part of acid and alkaline chemotaxis, respectively.^43,44^ In bacteria, dCACHE domain sensing is coupled to a variety of intracellular events, including activation of protein kinases/phosphatases and enzymes that turn over cyclic nucleotides. These sensory proteins classically function as homodimers, as suggested by the RDK1 model (Fig. 8b).

**Figure 8.**
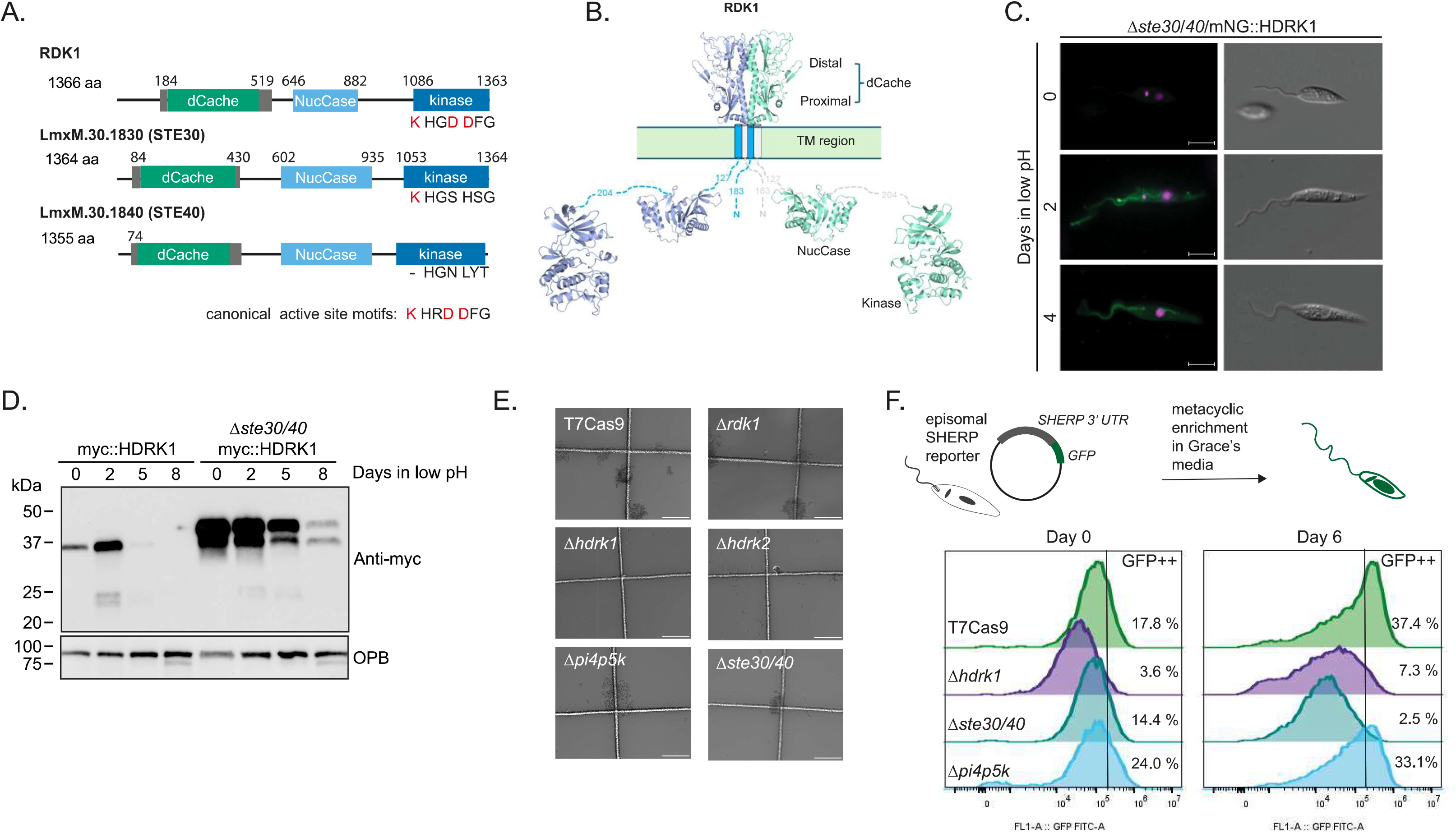
Transmembrane protein kinases are required for cell differentiation at low pH. A) Schematic of kinases with predicted transmembrane domains identified from the low-pH screen. Transmembrane domains are depicted as grey boxes. The dCACHE domain is shown in green, NucCase in light blue and the kinase domain in dark blue. Numbers indicate amino acid positions. The presence or absence of residues required for catalytic activity, K, D and D, is indicated under each kinase domain. Residues in red are required for kinase activity. B) Structural predictions for two RDK1 molecules forming a homodimer. Numbers indicate the number of residues between the modelled domains. C) Live-cell images showing mNG::HDRK1 in *Δste30/40* at day 0, 2 and 4 in low pH. Scale bar, 5 µm. D) Western blot of myc::HDRK1 in *bste30/40*. OPB was used as a loading control. E) Phase-contrast images showing attachment of T7Cas9 and mutant cell lines to gridded glass slides. Attached cell clumps appear as dark areas protruding from the + shape grid lines. Scale bar, 50 pm. F) Flow cytometry measuring SHERP expression in T7Cas9 and mutant lines at day 0 and 6 in low pH using a GFP reporter.

To determine whether STE30/40 acts upstream of HDRK1 in the low-pH sensing pathway, we tracked mNG::Hdrk1 localisation in *Δste30/40*. Signal at the basal body was weak at pH 7.4 but localised along the flagellum after two days in low pH, as seen in T7Cas9 cells, indicating a low-pH response (Fig. 8c). Western blotting of myc::HDRK1 demonstrated increased and prolonged HDRK1 expression in *Δste30/40* compared to T7Cas9, indicating a more complex relationship (Fig. 8d). Because the kinase mutants *hrdkl* and *Δpi4p5k* also exhibited morphological changes at low pH, we tested their capacity to adhere alongside *Δste30/40 and Δhdrk2*. Using gridded slides, we confirmed that all lines except *Δhdrk2* could adhere (Fig. 8e).

Phenotypic measurements were taken from *Δste30/40* at day 8 at low pH. Based on parameters by Sádlová et al.,^44^ *Δste30/40* lacked cells displaying metacyclic-stage morphology (Supplementary Fig. 9a). As SHERP is a known marker of the metacyclic promastigote stage,^45^ we expressed an episomal copy of a GFP-sherp reporter (pSSU-GFP-sherp) in the mut ants *Δhdrk1, Δste30/40* and *hpi4p5k*. Flow cytometry and fluorescent imaging showed an increased proportion of the population expressing high levels of GFP-SHERP by day 6 in T7Cas9 cells (Fig. 8f, Supplementary Fig 9b). This increased expression, linked to metacyclogenesis, was also observed in the *hpi4p5k* population but not in *Δhdrk1* or *Δste30/40*. Taken together, both HDRK1 and the LmxM.30.1830/40 locus are required for metacyclogenesis, but their relation to one another in responding to low pH remains to be determined.

## Discussion

Screening our kinome knockout library in defined medium reduced metabolic redundancy. Five-day pre-adaptation ensured that the screen identified acid-specific kinases rather than responses to nutrient shifts. To eliminate false-positive gains from population collapse, downstream characterisation of candidates was required, validating HDRK1 as acid-responsive and revealing interesting further phenotypes.

Our screen implicates phosphoinositide metabolism in environmental adaptation. Low-pH survival required PI4K, whereas PI4P5K loss conferred a survival advantage, indicating highly specific phosphoinositide balance requirements during acid stress. Transcriptomics and proteomics support this survival strategy. A PI4P5K-related protein (LmxM.20.0130) and myo-inositol-1-phosphate synthase (MIPS), the rate-limiting enzyme for *de novo* myo-inositol biosynthesis, were downregulated in *Δhdrk1* at low pH. MIPS downregulation restricts inositol production, limiting the phosphatidylinositol pool required for the lipophosphoglycan coat during metacyclogenesis. While our SHERP assay shows PI4P5K disruption alone does not impact metacyclogenesis, it is required for sand fly colonisation.^16^ Lipid balance importance is further reinforced by *Δhdrk2*, which contains a phosphoinositide-binding pleckstrin homology (PH) domain and produces a bulb structure at low pH.

Beyond metabolism, HDRK1 is a central developmental regulator. Although *Δhdrk1* cells adopt haptomonad-like morphology, they fail to attach to scratched plastic or the sand fly stomodeal valve. Attachment is a defining feature of the haptomonad stage and it is regulated by pH in the related kinetoplastids *Crithidia*^46^ and *Paratrypanosoma*^47^. Comparing our data with transcriptomic^5,47,48^ and proteomic datasets^32,46^ of attached cells revealed substantial molecular overlap despite the *Δhdrk1* attachment defect. This provides a unique tool to uncouple haptomonad differentiation from physical adhesion.

Our omics datasets reveal that *Δhdrk1* haptomonad-like cells upregulate attachment proteins SMP-1^34^ and KIAP2-4^32,49^ yet fail to adhere. Disrupted flagellar lipid homeostasis is known to cause membrane ballooning into an enlarged bulb over a shortened or coiled axoneme, coated by the lipid-anchored protein SMP-1.^50^ This closely mirrors our observations in *Δhdrk1* and *Δhdrk2* at low pH, corroborated by our omics data and Stains-All analysis showing lipid alterations. Flagellar membrane ballooning likely occurs due to disrupted protein-lipid interactions at the flagellum. The PH domain in HDRK2 points towards compromised flagellar lipid targeting, where failing to stabilise the membrane may deplete structural lipids and drive ballooning. Recent work links specific flagellar attachment zone (FAZ) proteins to adhesion. We observed decreased expression of FAZ7a and FAZ8, but no differential expression of FAZ2, FAZ5, or FAZ34, which are considered critical for attachment.^38,51^ Finally, cAMP has been associated with both pH taxis and attachment in parasites.^19,46^ We examined these pathways but detected no differential expression of PDE or CARP proteins. This absence of signalling alterations implies the defect is not a failure to sense attachment cues, but rather a structural inability to attach.

In our previous screen, HDRK1 mutants were enriched during metacyclic enrichment.^16^ This is because haptomonad-like cells have similar density to metacyclics, and a larger proportion of *Δhdrk1* cells become haptomonad-like than become metacyclic in a T7Cas9 cultures, leading to an artificial enrichment. Given that *Δhdrk1* cells fail to colonise the sand fly stomodeal valve, it was surprising to observe comparable haptomonad frequencies in *Δhdrk1*, T7Cas9 and the add-back (Fig. 3d), particularly as they were absent in KIAP null mutants.^32^ Instead, we observed fewer metacyclic cells and a large increase in elongated nectomonad-form parasites. One explanation is that *Δhdrk1* cells never reach the stomodeal valve and therefore do not encounter the lower-pH microenvironments necessary for further differentiation. Alternatively, physical attachment itself may trigger downstream developmental signalling. We propose that wild-type haptomonads utilise active AMPK pathways to reduce energy expenditure. At low pH, HDRK1 may restrain AMPK activation to allow commitment to the high-energy metacyclic form. In the absence of HDRK1, AMPK signalling persists, trapping the population in a quiescent, haptomonad-like state (Fig. 9). This model parallels *Trypanosoma brucei* biology, where AMPKα drives differentiation from the energy-intensive slender form to the quiescent, transmission-adapted stumpy form.^52^ While trypanosome stumpy forms are pre-adapted for the tsetse fly,^53^ *Leishmania* metacyclic forms are high-energy states pre-adapted for the mammal. Although the haptomonad stage may actively participate in mammalian infection,^5^ *Δhdrk1* cells show impaired infectivity.^16,54^ If HDRK1 functions as an AMPKα subunit, it seems paradoxical that its deletion leads to an “AMPK active” phenotypic state. This could be due to HDRK1 inhibiting the broader AMPK complex, or its absence triggering compensatory hypephosphorylation of alternative AMPK subunits. While *Δhdrk2* phenocopies *Δhdrk1*, their roles during mammalian infection diverge, as *Δhdrk2* infectivity was not impaired in the pooled screen.^16^

**Figure 9.**
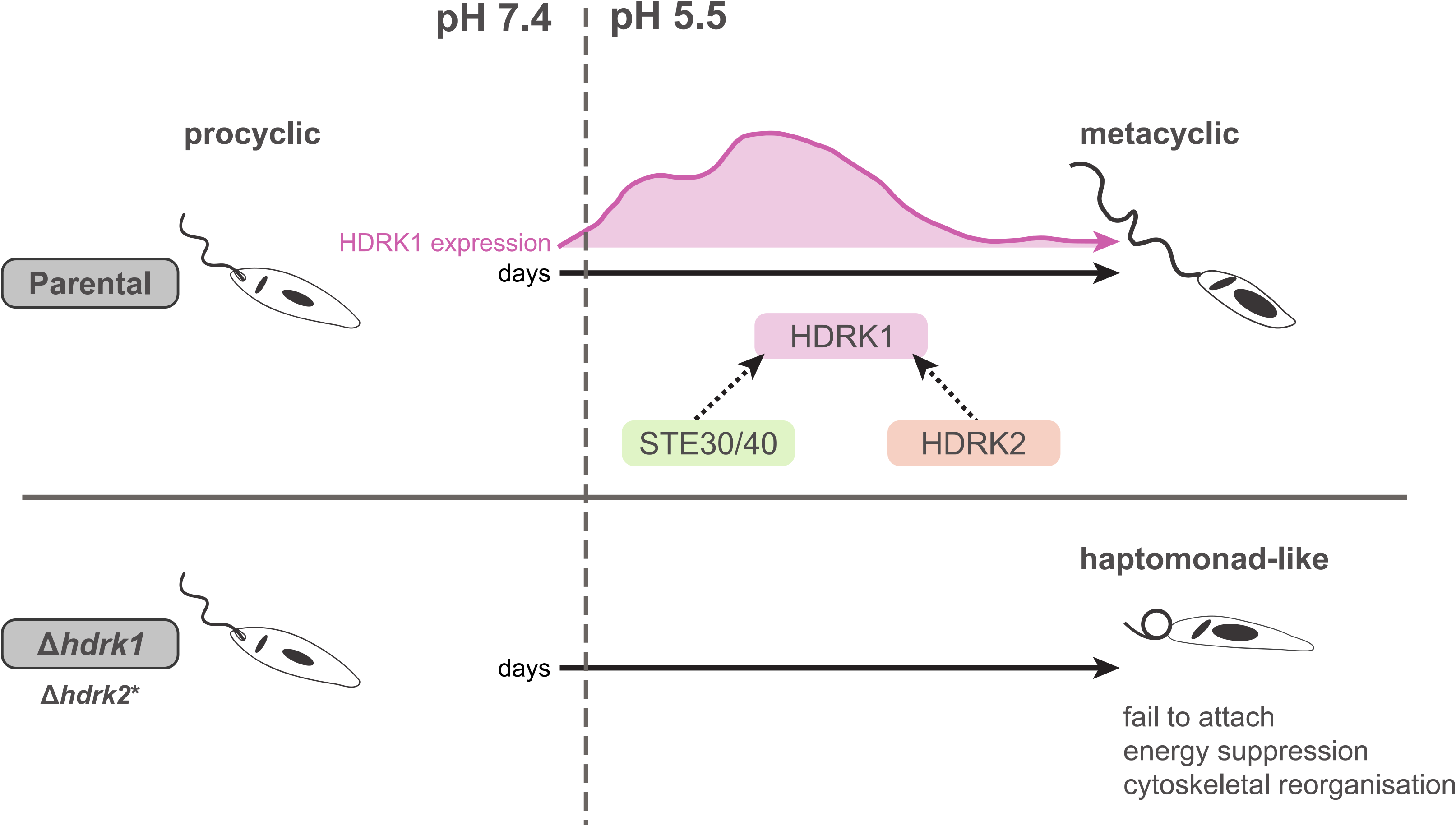
HDRK1 is required for metacyclic promastigote differentiation in response to low pH. Schematic illustrating that, in the parental cell line, exposure of *Leishmania* to low pH increases HDRK1 expression, leading to differentiation into metacyclic forms. STE30/40 and HDRK2 deletion affect HDRK1 expression and stability, suggesting they act within the HDRK1 pathway, although whether they function upstream or downstream of HDRK1 remains unknown, as indicated by dashed arrows. In the absence of HDRK1, cells become haptomonad-like cells at low pH, fail to attach *in vitro* and *in vivo*, and undergo energy suppression and cytoskeletal reorganisation. *HDRK2 deletion phenocopies the HDRK1 null mutant, with cells transforming into haptomonad-like cells and showing impaired cell adhesion *in vitro*.

How *Leishmania* senses environmental pH changes to trigger these kinase cascades remains an open question. While TOR1 is a known nutrient sensor important for promastigote differentiation,^55^ our datasets point to a distinct, TOR1-independent pH-sensing mechanism. Clues may lie among the other kinases identified in our low-pH screen. Only one, LmxM.25.1560, was upregulated in the HDRK1 omics data. However, putative environmental sensors would sit upstream of HDRK1, meaning differential expression is not required for them to share a signalling pathway. RDK1 and the two STE transmembrane kinases identified in our screen are predicted to possess extracytoplasmic dCACHE-type domains with striking structural similarity to the sensor domains of well-characterised bacterial two-component systems that sense and respond to changes in the extracellular environment. While previous studies localised these to the cytoplasm, N-terminal tagging may have disrupted their transmembrane insertion. Because *Leishmania* tightly regulates and stabilises intracellular pH,^56^ acidic pH is likely detected externally by surface-exposed sensors such as these STE kinases, rather than through internal cytoplasmic shifts.

Here we show that HDRK1 is integral to the acidic-pH stress response of *Leishmania*. HDRK1 is required for parasites to commit to the energy-intensive metacyclic form. While *Δhdrk1* cells fail to attach to surfaces, their adoption of haptomonad-like morphology provides a tool for decoupling the molecular biology required for this life cycle stage and the physical act of adhesion. Given that organisms frequently utilize multiple AMPK subunits to form tissue- or cue-specific complexes^52,57^ with multiple mechanisms of activation, HDRK1 highlights the complexity of parasite environmental sensing. Further research will be required to decipher the full pathway required for acidic-pH detection and signalling within *Leishmania*.

## Methods

### Cell culture

*Leishmania mexicana* (MNYC/BZ/62/M379) promastigotes expressing T7 RNA polymerase and Cas9^58^ were cultivated at 25°C in either HOMEM medium (Thermo Fisher Scientific) or M199 medium (Gibco, 15490574). Media were supplemented with 10% (v/v) heat-inactivated fetal calf serum (HIFCS; Gibco) and 1% penicillin/streptomycin (Sigma-Aldrich). Continuous selection was maintained with 50 µg/mL hygromycin (InvivoGen) and 75 µg/mL nourseothricin (Stratech Scientific Ltd). Procyclic promastigotes were subcultured at 1.5 x 10^5^ parasites/mL; cultures typically reached mid-log phase (5 x 10^6^ parasites/mL) by day 3 and stationary phase (2 x 10^7^ parasites/mL) by day 7. For differentiation assays, mid-log promastigotes (1.5 x 10^6^ parasites/mL) were washed twice in PBS and seeded into Grace’s insect medium (Sigma-Aldrich), adjusted to pH 5.5 and supplemented with 10% HIFCS, 1% penicillin/streptomycin, and 1% BME vitamins (Sigma-Aldrich). Flasks (non-vented, tissue-culture-treated; Corning) were laid flat and incubated at 25°C for up to 8 days.

### Pooled kinome-wide mutant library screen

Mid-log phase mutants from the kinome-wide barcoded library^16^ were grown separately and then pooled in equal proportions across four biological replicates, with each mutant added at 4 x 10^4^ cells/mL. Pooled cultures were grown in Nayak medium supplemented with 10% FCS and 1% penicillin/streptomycin, pH 7.4, at 25°C for 5 days to allow adaptation to the minimal medium. Samples for genomic DNA extraction were collected before adaptation and immediately after it (time 0). Following adaptation, cultures were washed twice in PBS and resuspended at 1 x 10^6^ cells/mL in fresh Nayak medium at pH 7.4 or adjusted to pH 5.5. After 48 h of incubation, genomic DNA was extracted, and barcode amplification and Illumina sequencing were carried out as previously described.^16^ Total barcode counts for each sample, including all biological replicates, are provided in Supplementary Table 1. To account for variation in sequencing depth, barcode counts for individual genes were normalised by expressing them as a percentage of the total read count within each sample (% representation). To quantify the impact of acidic conditions on individual mutants, a relative abundance ratio was calculated for each gene by dividing the normalised barcode representation at time 0, by the normalised barcode representation after 48 h incubation at either pH 5.5 or pH 7.4. Under this metric, lower ratio values indicate higher relative abundance of the corresponding mutant after 48 h, whereas higher ratio values indicate depletion. pH-specific effects were identified by subtracting the mean abundance ratio under acidic conditions (pH 5.5) from the mean abundance ratio under neutral conditions (pH 7.4). A positive difference in score therefore identifies genes whose disruption provided a relative survival advantage under pH 5.5. Statistical significance between the mean values was evaluated using multiple unpaired t-tests with Welch’s correction, with false discovery rates reported as q-values.

### Measurement of cell morphology

Mutants were grown individually to mid-log phase. Cells were fixed with formaldehyde at a final concentration of 1% (Alfa Aesar, 16% w/v, #43368) for 15 min. Fixed cells were washed in PBS by centrifugation at 1,000 x *g* for 1 min, and the pellet was gently resuspended in 20 µL PBS. A 10 µL aliquot of the suspension was added to a glass slide (Epredia SuperFrost Plus, #J1800AMNZ). Cells were imaged using an inverted Zeiss Axio Observer microscope with a 63x objective. Multiple fields of view were acquired for each cell line. Cell morphology measurements were performed in ImageJ 2.14 using the segmented line tool. For consistency, cell width was measured across the middle of the cell body, whereas flagellum length was measured from the flagellar tip to the edge of the cell body, following the curvature of the flagellum as closely as possible. For each cell line and time point, 50 cells were measured. Measurements were compared with T7Cas9 using a one-way ANOVA followed by Dunnett’s multiple comparisons test in GraphPad Prism. Measurements and ANOVA outputs are provided in Source Data. The percentage of cells with extra-axonemal material or a bulb was calculated from counting 100 cells across 10 images.

### CRISPR-Cas9 editing of *Leishmania*

CRISPR-Cas9-based editing was used to tag proteins, delete ORFs and to introduce precise codon edits within the HDRK1 ORF, resulting in defined amino acid substitutions in HDRK1. All oligonucleotide sequences and cell line identifiers are provided in Supplementary Information. Tagging and knockout edits were designed using the LeishGEdit toolbox.^58^ DNA templates for genetic edits were electroporated into *Leishmania* using an Amaxa 4D transfection kit and Amaxa 4D electroporation machine. After recovery, cells were selected with 50 µg/mL puromycin, 10 µg/mL blasticidin or 25 µg/mL G418 (InvivoGen).

Precision edits were designed manually and carried out using an approach analogous to that described by McNiven^30^. Briefly, two overlapping primers were designed to generate a 150—160 bp repair cassette containing the desired codon substitutions, with more than 50% of codons changed.

Following transfection, cells were allowed to recover for 2-12 h in M199 medium supplemented with 20% FCS and 10 µM 6-biopterin before cloning into 96-well plates in the same supplemented medium. Diagnostic PCRs were used to identify clones in which both alleles had been edited, and edits were confirmed by Sanger sequencing (Supplementary Fig. 3).

### HDRK1 add-back and reporter plasmid transfection

An exogenous copy of HDRK1 was reintroduced into the HDRK1 null line by cloning the HDRK1 ORF into the pRIB transfection vector pGL2398. *Xhol* and *Notl* were used to excise the CRK3 gene, which was replaced with the HDRK1 ORF. For transfection, 10 μg of DNA was prepared by digestion with *PacI* and *PmeI*, separated by gel electrophoresis, and extracted using a Qiagen Gel DNA extraction kit. The resulting DNA was precipitated with ethanol, washed and resuspended in sterile H_2_O.

A pSSU-GFP-sherp reporter plasmid^59^ was modified by replacing the hygromycin selectable marker with a neomycin selectable marker amplified from pTNeo_V1.^58^ 10 μg of purified vector was transfected into mid-log phase *Leishmania*, and cells were selected with 25 µg/mL G418 (InvivoGen) for episomal expression of the reporter.

### Flow cytometry

For cell cycle analysis, 5 x 10^6^ cells were washed in PBS supplemented with 5 mM EDTA. The resulting pellet was resuspended in 70% methanol prepared in PBS containing 5 mM EDTA and fixed overnight at 4°C. Cells were washed twice in PBS containing 5 mM EDTA by centrifugation at 1,000 x *g* for 5 min and resuspended in staining solution containing 10 µg/mL propidium iodide (PI) and 10 µg/mL RNase A. Samples were incubated at 37°C for 45 min and transferred to a 96-well plate. For each cell line, 10,000 events were acquired on a CytoFLEX S flow cytometer.

GFP signal in T7Cas9 and mutant lines containing episomal copies of the GFP-SHERP reporter was measured using live-cell flow cytometry. PI (0.5 µg/mL final concentration) was added 10 min before data acquisition as a viability stain. Intact parasites were gated using forward scatter area (FSC-A) versus side scatter area (SSC-A), and single cells were selected using FSC-H versus FSC-A doublet discrimination. Viable cells were gated using PI exclusion (PI vs. FSC-A), using a heat-treated control to define the PI-positive population. GFP-SHERP expression levels were then compared between samples.

### Sand fly infections

The colony of *Lutzomyia longipalpis*, originating in Jacobina, Brazil, was maintained in the insectary of the Department of Parasitology, Charles University, Prague, under standard conditions: 26°C, access to 50% sucrose, 60-70% humidity and a 14 h light/10 h dark photoperiod, as described previously.^60^

Promastigotes from log-phase cultures (day 3–4 in culture) were washed twice in saline and resuspended in heat-inactivated ram blood (LabMediaServis) at a concentration of 1 x 10^6^ promastigotes/mL. Female sand flies, 5–9 days old, were infected by feeding through a chick-skin membrane (BIOPHARM) on the promastigote-containing blood suspension. Engorged sand flies were maintained under the same conditions as the colony. The location of *Leishmania* infections in the sand fly digestive tract (endoperitrophic space, abdominal midgut, thoracic midgut, cardia region and colonisation of the stomodeal valve) was determined by dissection and examination by light microscopy on days 1, 4–5 and 9 post blood meal (PBM). Parasite loads were graded according to Myskova et al.,^61^ as light (<100 parasites per gut), moderate (10–1,000 parasites per gut) or heavy (>1,000 parasites per gut). Sand fly infection experiments were performed twice.

### Morphometry of parasites dissected from sand fries

Gut smears of L. mexicana-infected females at 9 days PBM were fixed with methanol, stained with Giemsa and examined by light microscopy using an oil-immersion objective. For each *Leishmania* line, 300 randomly selected promastigotes from six sand flies/smears were measured, with three smears analysed per each experimental infection. Parasite body length, flagellar length and body width were measured using ImageJ software. Three morphological forms were distinguished: elongated nectomonads, body length >14 µm; metacyclic promastigotes, body length ≤ 14 µm and flagellar length >2 times body length; and leptomonads (short promastigotes), body length ≤ 14 µm and flagellar length < 2 times body length.^44^ In addition, haptomonads with a very short broad flagellum were evaluated according to Walters.^31^ Differences in cell body or flagellum measurements were tested by independent samPles t-tests and differences in representation of morphological forms by chi-square test. All the analyses were performed using SPSS version 23.

### Attachment assays

For attachment to scratched plastic, Melinex film (Agar Scientific) was scratched with P2000 ultra-fine sandpaper, cut into 1 cm x 5 cm strips, immersed in 70% ethanol, agitated and air-dried. Mid-log parasites were seeded at 1.5 x 10^6^/mL in Grace’s media in 75 cm^2^ flasks containing the scratched plastic strips. Flasks were placed on their sides and incubated for 5 days. The plastic strips were then removed with tweezers, washed twice by gentle agitation in PBS in a petri dish for 5 min, stained with Giemsa (Sigma 48900) and washed again. Plastic strips were imaged at 1x magnification using a ChemiDoc MP imaging system (Bio-Rad) and at 40x magnification using an inverted light microscope. For attachment to gridded glass slides, cells were grown in 12-well plates containing gridded glass coverslips (Grid-500, IB-10816, Thistle Scientific) as described previously.^62^ Briefly, cells were seeded at 1 x 10^6^ cells/mL into the wells containing coverslips, incubated for 48 h in HOMEM medium and washed gently before addition of 50 μL CyGel and a glass slide.

### Live-cell microscopy

mNeonGreen-tagged proteins were examined in live cells. 1 x 10^6^ mid-log cells were harvested by centrifugation at 1,200 x *g* for 10 min and washed in PBS before incubation with 5 µg/mL Hoechst 33342 at room temperature for 10 min. Excess Hoechst was removed by washing the cells in 10 mL of ice-cold PBS, followed by centrifugation at 1,200 x *g* for 10 min. The supernatant was removed completely, cells were resuspended in 50 μL CyGel on ice and 25 μL of cell suspension was dispensed onto a slide. Cells were covered with a coverslip and imaged immediately. SHERP expression was imaged in live cells settled directly onto positively charged glass slides, as described previously. Briefly, cells were harvested by centrifugation at 1,200 x *g* for 7 min, washed in PBS and incubated with 1 µg/mL Hoechst 33342 at room temperature for 10 min. Cells were then harvested by centrifugation, resuspended in 10−300 µL PBS, and 5 µL of the cell suspension was dispensed onto a slide and imaged.

### Fixed-cell microscopy

Mid-log phase cells were washed in PBS and fixed in 2% paraformaldehyde for 15 min at room temperature. Cells were washed twice in PBS by centrifugation at 1,000 x *g* for 10 min. The final pellet was gently resuspended in approximately 20−50 µL PBS, and 5 µL of cell suspension was added to a slide before imaging.

### Immunofluorescence

For immunolocalisation of ISP1, cells were washed in PBS and allowed to adhere to poly-L-lysine coated coverslips for 15 min at 37°C. Excess solution was removed, and adhered cells were washed once in PBS before fixation with 4% PFA for 15 min at room temperature. Residual PFA was quenched with 0.1 M Tris-HCl (pH 8.5) for 5 min. Cells were permeabilised with 0.5% Triton X-100 in PBS for 10 min and subsequently blocked with 5% BSA and 0.01% saponin in PBS for 1 h at RT. Primary antibody incubation was performed for 2 h at RT in a humidified chamber using sheep anti-ISP1 (1:25 dilution in PBS containing 5% FBS).^33^ Following three washes with PBS, cells were incubated for 1 h at RT with Alexa Fluor 488-conjugated donkey anti-sheep secondary antibody (1:2000 dilution, Thermo Fisher). Samples were washed three times in PBS, counterstained with 10 µg/mL DAPI in H_2_O, and subsequently washed in PBS and H_2_O. Coverslips were mounted onto glass slides using ProLong Diamond Antifade Mountant (Thermo Fisher) before imaging on a Zeiss Axio Observer inverted microscope with 63x oil-immersion objective lens.

### Transmission Electron Microscopy and Scanning Electron Microscopy

For transmission electron microscopy (TEM), mid-log phase cells were fixed for 10 min in a fixative containing 8% formaldehyde and 5% glutaraldehyde in 100 mM sodium phosphate buffer, pH 7.3, mixed 1:1 with culture medium. Cells were then fixed for a further 30 min in 4% formaldehyde and 2.5% glutaraldehyde in 100 mM sodium phosphate, pH 7.3. Subsequent washes in phosphate buffer, ethanol and acetone were carried out as previously described.^63^

For scanning electron microscopy (SEM), 2 x 10^7^ cells were washed three times in serum-free medium and once in PBS before resuspension in 100 µL PBS and settling onto coverslips. Coverslips were washed once in 0.1 M sodium cacodylate buffer before fixation with 1 mL of 2.5% glutaraldehyde in 0.1 M sodium cacodylate buffer. Samples were then fixed in 1% tannic acid in 0.1 M sodium cacodylate buffer and dehydrated through graded ethanol series, 25−100%, for 15 min at each step. Ethanol was exchanged with hexamethyldisilazane (HMDS) through two changes, and samples were left to dry in a desiccator overnight. Samples were mounted onto SEM stubs, sputter-coated with 5 nm of gold/palladium using a Polaron SC7640 sputter coater, and imaged using a JEOL JSM-7800F scanning electron microscope operating at and an accelerating voltage of 8 kV.

### Protein gels and western blotting

For cell lysis, parasites were resuspended in RIPA buffer (0.1% sodium dodecyl sulfate, 0.5% sodium deoxycholate, 1% IGEPAL CA-630, 0.1 mM EDTA, 125 mM NaCI, 50 mM Tris-HCl pH 7.5, and 1 mM sodium orthovanadate) containing 2x HALT Protease Inhibitor Cocktail (Thermo Fisher Scientific) and 2x PhosSTOP phosphatase inhibitor (Sigma). BaseMuncher Endonuclease (Abcam), 1 µL, was added to each sample, and cells were sonicated at 4°C for three cycles of 30 s on and 30 s off using a Bioruptor Sonication system (Diagenode). Protein concentration in cell lysates were quantified using the Bradford assay, and 23 µg of protein was used per sample. NuPAGE sample buffer supplemented with 50 mM dithiothreitol (DTT) was added to each sample and loaded onto 4−12% Bio-Rad Mini-PROTEAN stain-free protein gels. Gels were run in 1x Tris/glycine/SDS PAGE running buffer and transferred to nitrocellulose membranes using the iBlot system (Invitrogen). Membranes were blocked with 5% skimmed milk in Tris-buffered saline with Tween 20 (TBS-T) for lh before incubation with mouse monoclonal anti-myc antibody, clone 4A6, 1:3,000 (Millipore), diluted in 5% skimmed milk in TBS-T for 1 h at room temperature. After washing, membranes were incubated with horseradish peroxidase (HRP)-conjugated anti-mouse IgG (H+L) secondary antibody, 1:20,000 (Molecular Probes), in 5% skimmed milk in TBS-T for 1 h at room temperature. Membranes were washed, incubated with Clarity Max Substrate (Bio-Rad) and developed using a Chemidoc MP imaging system (Bio-Rad). For relative expression quantification, membranes were stripped using Restore Western Blot Stripping Buffer according to the manufacturer’s instructions, blocked again, and reprobed with sheep anti-OPB antibody, 1:20,000, diluted in 5% skimmed milk in TBS-T for 1h at room temperature. After washing, membranes were incubated with HRP-conjugated anti-sheep IgG secondary antibody, 1:5,000 (antibodies.com), under the same conditions and developed as described above. Band intensities of myc-tagged proteins were normalised to the corresponding OPB band intensity using ImageJ.

For Stains-All staining, samples were separated on NuPAGE gels. Gels were equilibrated in 30% ethanol in water for 1 h at room temperature, incubated overnight in working Stains-All solution protected from light, and washed with water until the gels no longer floated. Gels were imaged using a ChemiDoc MP imaging system (Bio-Rad).

### RNA-seq

Total RNA was extracted from triplicate cultures of T7Cas9, *Δhdrk1* cl7 and *Δhdrk1* cl8 grown in Grace’s media on days 0, 4 and 7, using the NEB Monarch Total RNA Miniprep kit. Libraries were sequenced by Novogene,Cambridge, UK, using 150 bp paired-end reads. Raw reads were assessed using FastQC^64^ and MultiQC^65^ with default parameters. HISAT2 version 2.2.1^66^ was used with default parameters to map reads onto the L.mex T7Cas9 genome. Picard Tools version 2.24.0 was used for post alignment sorting and HTSeq-count version 0.11.1 was used to count alignments against features using the union mode.

Differential expression analysis and subsequent principal component analysis (PCA) were performed on the resulting counts using the DESeq2 R package^67^. Differential expression was assessed by comparing T7Cas9 with *Δhdrk1* cl7 and *Δhdrk1* cl8 at each of the three time points (day 1, 5 and 8) and using replicate, time and type as factors in the model design. A 5% false discovery rate (FDR) threshold was used for significance. Outputs of less than 0.05 significance were filtered and a variance shrinking package - apeglm^68^ was used to reduce background noise in the data. PCA analysis was done using the plotPCA function within DESeq2 and by rlog transforming the counts. Outputs were then filtered further to contain genes differentially expressed in both *Δhdrk1* cl7 and *Δhdrk1* cl8. The heatmap generating tool, ClustVis^69^ was used to cluster the datasets according to similar groupings. GO term enrichments were generated for the DE expressed genes using the TriTrypDB and REVIGO websites.

### Proteomics

For each sample, 6-8 x 10^8^ cells were harvested, washed twice in PBS and resuspended in 500 µL lysis buffer containing 5% SDS and 50 mM TEAB, pH 8.5. Samples were probe-sonicated on ice for two 10 s cycles at 60% amplitude, followed by additional lysis using a Bioruptor (Diagenode) for four cycles, 30 s ON, 30 s OFF. Samples were centrifuged at 20,000 x *g* for 15 min at room temperature before supernatants were collected. Samples were diluted 1:1 with aqueous 5% (w/v) SDS in 50 mM triethylammonium bicarbonate (TEAB). Proteins were reduced with 5.7 mM dithiothreitol by heating to 55°C for 15 min, before alkylation with 22.7 mM iodoacetamide at room temperature for 10 min. Protein samples were acidified by addition of 6.5 μL of aqueous 27.5% (v/v) phosphoric acid, then precipitated by seven-fold dilution into 100 mM TEAB 90% (v/v) methanol. Precipitated protein were captured on S-Traps (ProtiFi − C0-micro) and washed five times with 165 µL 100 mM TEAB in 90% (v/v) methanol before digestion with 20 µL 0.1 µg/µL Promega Trypsin/Lys-C mix (V5071) in aqueous 50 mM TEAB at 47°C on a hot plate for 2 h. Peptides were recovered from S-Traps by centrifugation at 4,000 x *g* for 60 s. S-Traps were washed with 40 µL aqueous 0.2% (v/v) formic acid and 40 µL 50% (v/v) acetonitrile in water, and washes were combined with the first peptide elutions. Peptide solutions were dried in a vacuum concentrator then resuspended in 50 µL aqueous 50 mM TEAB for TMT labelling. Peptides were labelled using 10-Plex TMT reagents (Thermo Fisher Scientific, 90110), following the manufacturer’s procedure. Haptomonad-like samples (day 7) were labelled with TMT reagents 126, 127N, 127C, 128N and 128C. Procyclic promastigote samples (day 0) were labelled with TMT reagents 129N, 129C, 130N, 130C and 131. Samples were combined after labelling, dried in a vacuum concentrator and resuspended in 100 μL water for fractionation by high-pH reversed phase C18 HPLC.

Samples were loaded onto an Agilent 1260 II HPLC system equipped with a Waters XBridge 3.5 µm C18 column (2.1 mm x 150 mm, Waters). Separation used gradient elution with two solvents: solvent A, aqueous 0.1% (v/v) ammonium hydroxide; solvent B, acetonitrile containing 0.1% (v/v) ammonium hydroxide. The flow rate was 200 µL/min and the column temperature was 40°C. The linear multi-step gradient profile for the elution was 5-35% B over 20 min, 35-80% B over 5 min, followed by washing with 80% solvent B for 5 min before returning to initial conditions and re-equilibrating for 7 min prior to subsequent injections. Eluate was collected at 1 min intervals into LoBind Eppendorf tubes. Peptide elution was monitored by UV absorbance at 215 and 280 nm. Fractions were taken by the minute from 17-27 min, generating 10 fractions. Fractions corresponding to 14-17 min were pooled, as were fractions corresponding to 27-30 min, giving a total of 12 fractions for LC-MS. Peptide fractions were dried in a vacuum concentrator before reconstitution in 20 µL aqueous 0.1% (v/v) trifluoroacetic acid (TFA).

Fractionated TMT-labelled peptides were loaded onto an M-Class nanoflow UPLC system (Waters) equipped with a nanoEase M/Z Symmetry 100 Å C18, 5 µm trap column (180 µm x 20 mm, Waters) and a PepMap 2 µm, 100 Å, C18 EasyNano nanocapillary column (75 µm x 500 mm, Thermo Fisher Scientific). The trap wash solvent was aqueous 0.05% (v/v) trifluoroacetic acid, and the trapping flow rate was 15 µL/min. The trap was washed for 5 min before switching flow to the capillary column. Separation used gradient elution of two solvents: solvent A, aqueous 0.1% (v/v) formic acid; solvent B, acetonitrile containing 0.1% (v/v) formic acid. The flow rate for the capillary column was 300 nL/min, and the column temperature was 40°C. The linear multi-step gradient profile was 3-10% B over 7 min, 10-35% B over 80 min and 35-99% B over 10 min, followed by washing with 99% solvent B for 8 min. The column was returned to initial conditions and re-equilibrated for 15 min before subsequent injections.

The nanoLC system was interfaced to an Orbitrap Fusion hybrid mass spectrometer (Thermo Fisher Scientific) with an EasyNano ionisation source (Thermo Fisher Scientific). Positive ESI-MS, MS2 and MS3 spectra were acquired using Xcalibur software, version 4.0 (Thermo Fisher Scientific). Instrument source settings were: ion spray voltage, 2,100 V; sweep gas, 0 Arb; and ion transfer tube temperature; 275°C. MS1 spectra were acquired in the Orbitrap with: 120,000 resolution, scan range *m/z* 380-1,500, AGC target, 2 x 10^5^ and maximum fill time 50 ms. Data-dependent acquisition was performed in topspeed mode using a 4 s cycle, selecting the most intense precursors with charge states 2-6. Dynamic exclusion was performed for 50 s after precursor selection, and the minimum threshold for fragmentation was set at 3 x 10^4^. MS2 spectra were acquired in the linear ion trap with scan rate set to turbo, quadrupole isolation of 1.2 *m/z*, CID activation, activation energy of 35%, AGC target 1x 10^4^, first mass m/z 120 and maximum fill time 35 ms. MS3 spectra were acquired in multi notch synchronous precursor selection mode (SPS-MS3), selecting the five most intense MS2 fragment ions between m/z 400 and 1,000. SPS3 spectra were measured in the Orbitrap mass analyser using 50,000 resolution, quadrupole isolation of 1 *m/z*, HCD activation type, collision energy of 65%, scan range m/z 110-500, AGC target 4 x 10^5^ and maximum fill time 10 ms. Acquisitions were arranged by Xcalibur to inject ions for all available parallelizable time.

Peak lists in .raw format were imported into PEAKS StudioX Pro, version 10.6 (Bioinformatics Solutions Inc.), for peak picking, database searching and relative quantification. MS2 peak lists were searched against LmexCas9T7-prot.fasta appended with common proteomic contaminants. Search criteria were specified as follows: enzyme, trypsin; maximum missed cleavages, 1; fixed modifications, TMT10plex on lysine residues and peptide N-termini, and carbamidomethylation on cysteine residues; variable modification, oxidation of methionine; peptide tolerance, 3 ppm; and MS/MS tolerance, 0.5 Da. Peptide identifications were filtered to achieve a 1% peptide-spectrum match false discovery rate, as assessed empirically against a reversed database search. Protein identifications were further filtered to require a minimum of two unique peptides per protein. TMT reporter ion intensities, used as measures of relative inter-sample peptide abundance, were extracted from MS3 spectra for quantitative comparison. For significance testing, a pairwise ANOVA was conducted between the two groups. A total of 3,833 proteins met the quantification thresholds. After multiple testing correction of ANOVA-derived significance values in PEAKS, quantification significance scores >18.58 and >28.02 were considered significant at <5% FDR and at <1% FDR, respectively, for discrimination between groups.

### HDRK1 protein expression and ADP-Glo assay

The HDRK1 ORF was codon-optimised and synthesised by Aruru Molecular Ltd. The ORF was PCR-amplified using oligonucleotides listed in Supplementary Information and cloned into an expression plasmid containing a maltose-binding protein (MBP) tag and a His tag. The plasmid was transformed into BL21(DE3) *E. coli*. Protein expression was induced using auto-induction media at 20°C overnight. Ni^2+^-affinity chromatography was used to purify N-His-MBP-HDRK1 from crude protein extract. Following application to the Ni^2+^-affinity column, bound protein was washed and then subsequently eluted using 30−500 mM imidazole gradient. Fractions containing eluted protein were analysed by SDS-PAGE to confirm purity. Protein fractions from Ni^2+^-affinity chromatography were pooled and placed into a dialysis bag containing H3C-protease and then dialysed overnight to cleave the MBP tag and remove imidazole. Analysis of pre- and post-digestion samples confirmed successful protease cleavage. The digestion products were recovered following dialysis and further purification was performed to remove the N-His-MBP tag and H3C-protease, which was also His-tagged. Following application to Ni^2+^-affinity chromatography, the column was washed first with a buffer containing 0 mM imidazole and then with a buffer containing 30 mM imidazole. HDRK1 protein eluted during the second imidazole wash. Further bound protein was subsequently eluted using an elution buffer containing 20 mM sodium phosphate pH 7.4, 300 mM NaCl and a 30-500 mM imidazole gradient. Fractions containing eluted HDRK1 protein were analysed by SDS-PAGE to confirm purity and product size. Fractions containing HDRK1 protein were pooled to ensure that the final concentration remained above 0.5 mg/mL. Protein concentration was estimated using data from the ÄKTA pure software and confirmed by absorbance at UV280nm using a spectrophotometer together with a predicted extinction coefficient. Protein was aliquoted into 1 mL aliquots for storage and subsequent use.

HDRK1 kinase activity was tested against a peptide panel from MRC PPU (University of Dundee), detailed in Supplementary Fig. 2, using the ADP-Glo assay (Promega) according to kit instructions.

### Structural predictions

The HDRK1 AlphaFold-predicted protein structure and human AAPK2 protein structure (UniProt: P54646) were aligned using the RCSB Pairwise Structural Alignment tool (https://www.rcsb.org/alignment). The protein sequences for these genes were aligned using MultAlin.^70^

## Data repositories

The raw sequencing files (FASTQ) for the and RNA-seq experiments, along with associated metadata, have been deposited in the NCBI BioProject database under the accession code PRJNA1499433. Read counts for the Bar-seq experiment are provided in Supplementary Table 2. Mass spectrometry raw files and proteomic results are deposited in the MassIVE database under accession code MSV000102504 and cross-referenced in ProteomeXchange under accession code PXD081187. Processed protein quantification data can be found in Supplementary Table 4.

## Code availability

The code used for Bar-seq data processing was previously described.^16^ All sequencing mapping, mass spectrometry database searches, and downstream quantitative analyses were performed using standard software tools as described in the Methods section.

## Additional information

Supplementary Information: List of oligos, Cell lines, cell morphology (Fig. 1d)

Supplementary Table 1: Bar-seq counts

Supplementary Table 2: RNA-seq raw counts

Supplementary Table 3: RNA-Seq DE analysis

Supplementary Table 4: Proteomic raw counts

Supplementary Table 5: Proteomic DE

## Acknowledgments

We thank the Bioscience Technology Facility at the University of York for technical support, including Meg Stark for transmission and scanning electron microscopy, Karen Hogg and Sukhveer Mann for flow cytometry, Sally James and Lesley Gilbert for Illumina sequencing services, Katherine Newling for sequencing barcode retrieval, Rebecca Preece for protein expression of HDRK1 and Alastair Droop for depositing sequencing and RNA-seq data to a public repository. We also thank Pegine Walrad for the SHERP reporter plasmid.

## Grants and Funding

This work was supported by a Wellcome Investigator award (223045/Z/21/Z) and Wellcome Trust Career Development award (226513/Z/22/Z). The York Centre of Excellence in Mass Spectrometry was created thanks to a major capital investment through Science City York, supported by Yorkshire Forward with funds from the Northern Way Initiative, and subsequent support from EPSRC (EP/K039660/1; EP/M028127/1). Sand fly investigations were supported by the Czech Science Foundation (GACR 25-15318S).

## Author information

### Author contributions

N.B. and J.C.M. conceived the study. J.C.M. supervised N.B., and N.B. secured independent funding and supervised L.V.L. N.B., L.V.L., J.A.B. and C.M. carried out lab experiments. J.S., B.B. and P.V. conducted sand fly investigations. S.F. carried out quality control, mapping and RNA-seq analysis. A.D. and C.T. designed and executed proteomics, including protein samples labelling, mass spectrometry and peptide mapping. J.A.B., N.B. and A.J.W. carried out structural bioinformatics. A.J.W. produced structural models. N.B. and L.V.L prepared the manuscript, A.J.W. and J.C.M. made substantial edits. All authors reviewed, edited and approved the final manuscript. J.C.M. and N.B. acquired the funding.

## Supplementary figures

**Supplementary figure 1.**
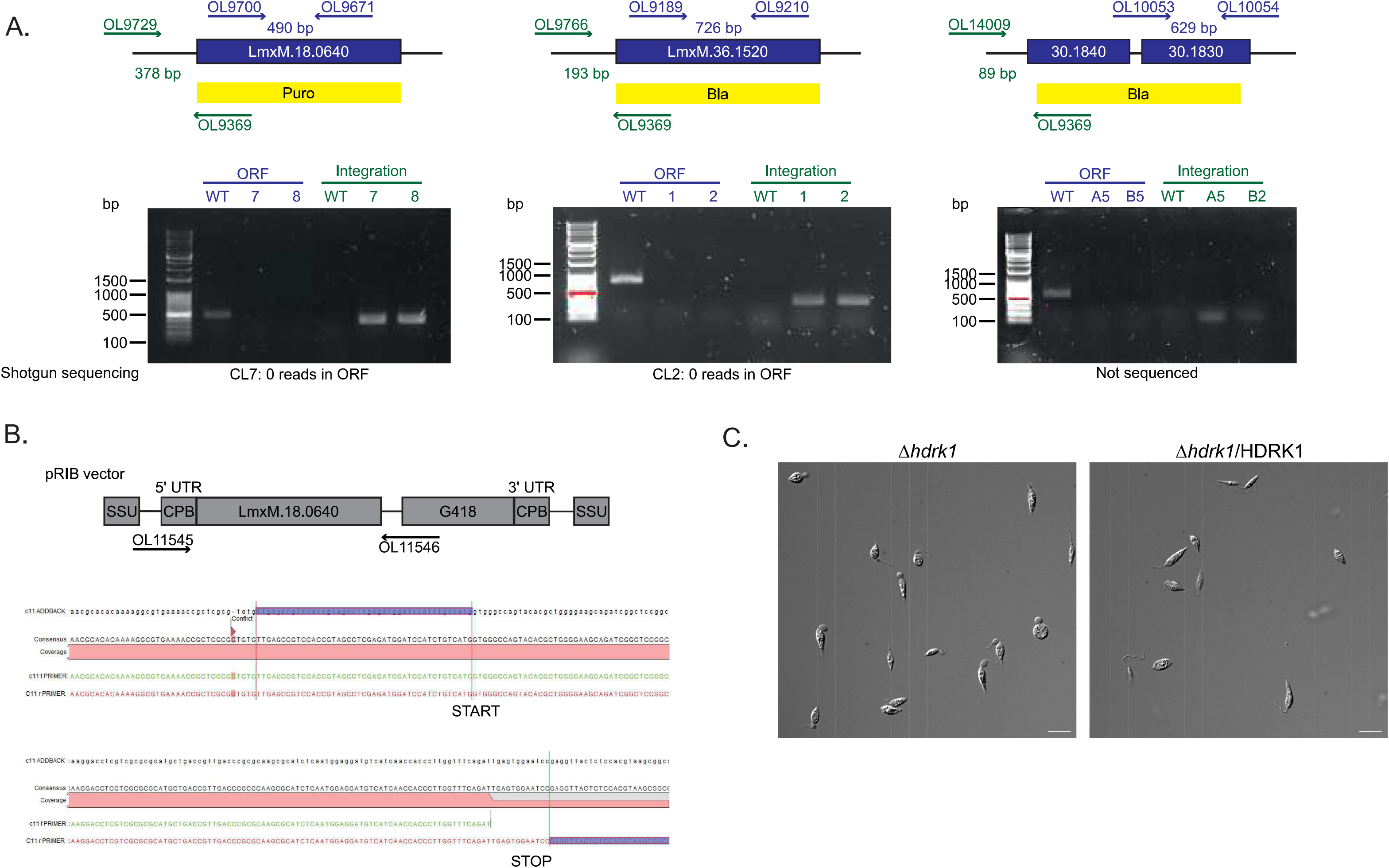
Generation and validation of cell lines. A) Schematics showing primers used for diagnostic PCRs for the *Δhdrk1, Δhdrk2* and *Δste30/40* null mutants (LmxM.18.0640, LmxM.36.1520 and LmxM.30.1830/40) and expected product sizes. Primers shown in blue amplify a region within the open reading frame (ORF), which is present in T7Cas9 but absent in the null mutants. Integration primers, shown in green, were designed upstream of the targeted locus, with a common reverse primer located within the repair cassette. The agarose gel shows absence of this product in T7Cas9 and presence in the null mutants. B) Schematic of the pRIB vector and cladogram showing sequencing-based confirmation of correct *hdrk1* insertion into the vector. C) Phase-contrast images of *Δhdrk1* and add-back after 8 days at low pH. Scale bar, 10 μm.

**Supplementary figure 2.**
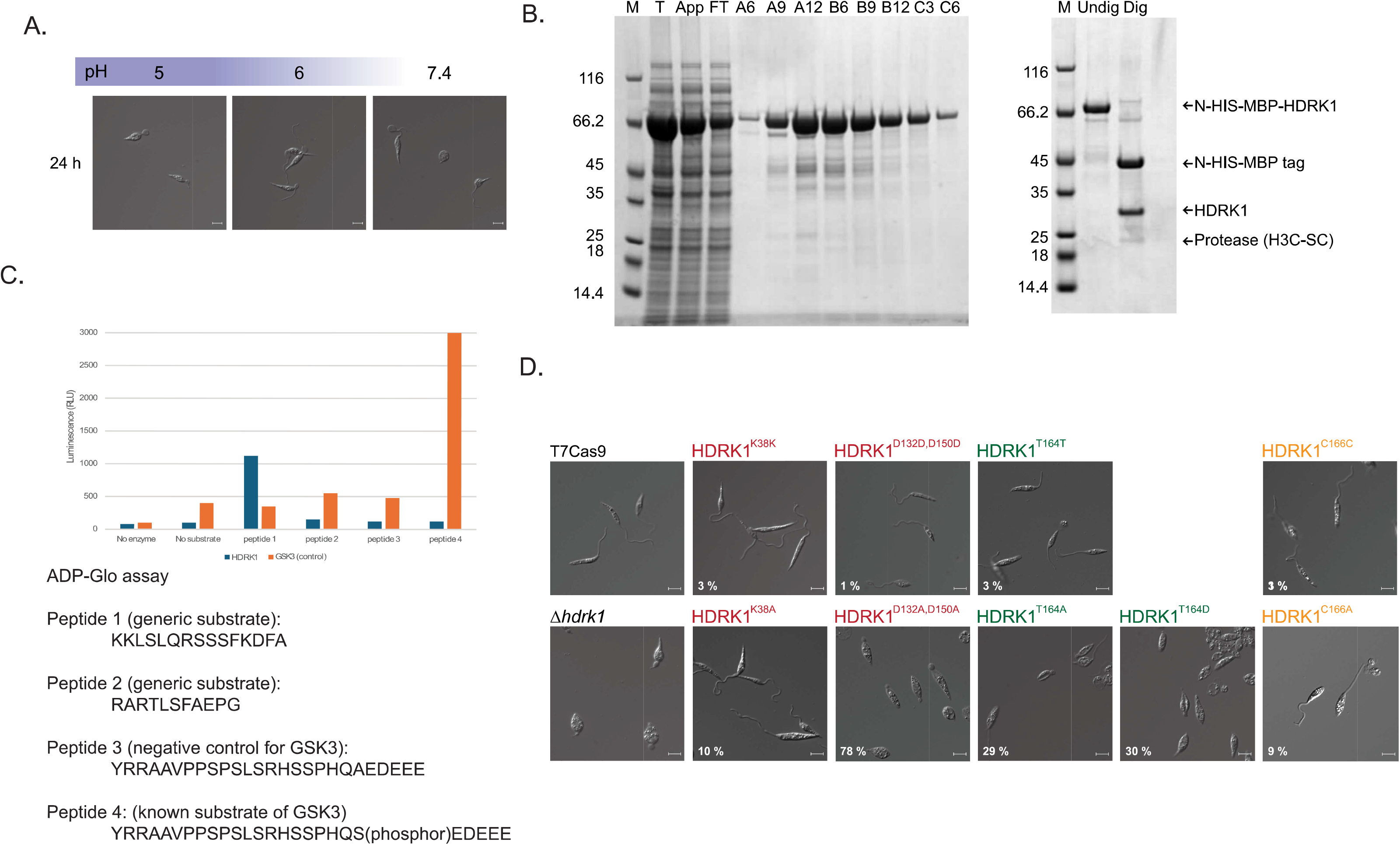
A) Phase-contrast images *of Δhdrk1* cells grown for 7 days in Grace’s media, pH 5.5, before washing and transfer to HOMEM medium at pH 5.0, 6.0 or 7.4 for 24 h. B) Expression of N-HIS-MBP-HDRK1 shown on SDS-PAGE gel. Digestion lead to cleavage of the N-HIS-MBP tag from HDRK1. C) ADP-Glo assay using recombinant HDRK1 protein. Peptides 1-4 were purchased from the University of Dundee. GSK3 protein was used as a positive control with known activity against peptide 4. D) Phase-contrast images of HDRK1 precision-edit mutants after 7 days at low pH. Scale bar, 5 µm. White text shows the percentage of cells in each population with extra-axonemal material.

**Supplementary figure 3.**
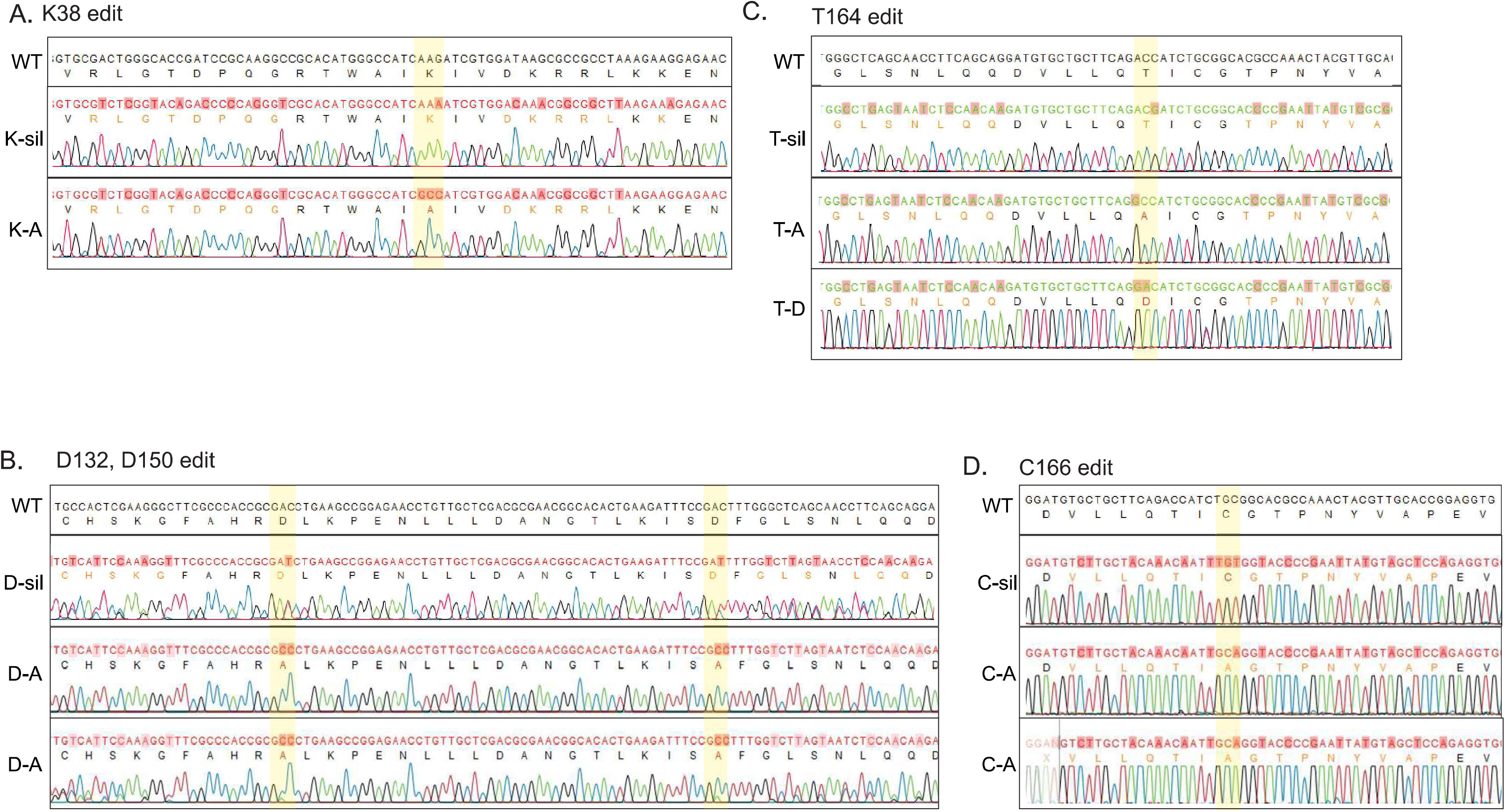
Sanger sequencing confirming precision edits in HDRK1. A) Editing of lysine at position 38 from AAG to a synonymous codon, AAA (silent mutant) or to alanine, GCC. B) Editing of aspartic acid residues at position 132 and 150 from GAC to synonymous codon, GAT (silent mutant), or to alanine, GCC. C) Editing of threonine at position 164 from ACC to the synonymous codon, ACG (silent mutant), alanine, GCC, or aspartic acid, GAC. D) Editing of cysteine at position 166 from TGC to the synonymous codon, TGT (silent mutant), or alanine, GCA.

**Supplementary figure 4.**
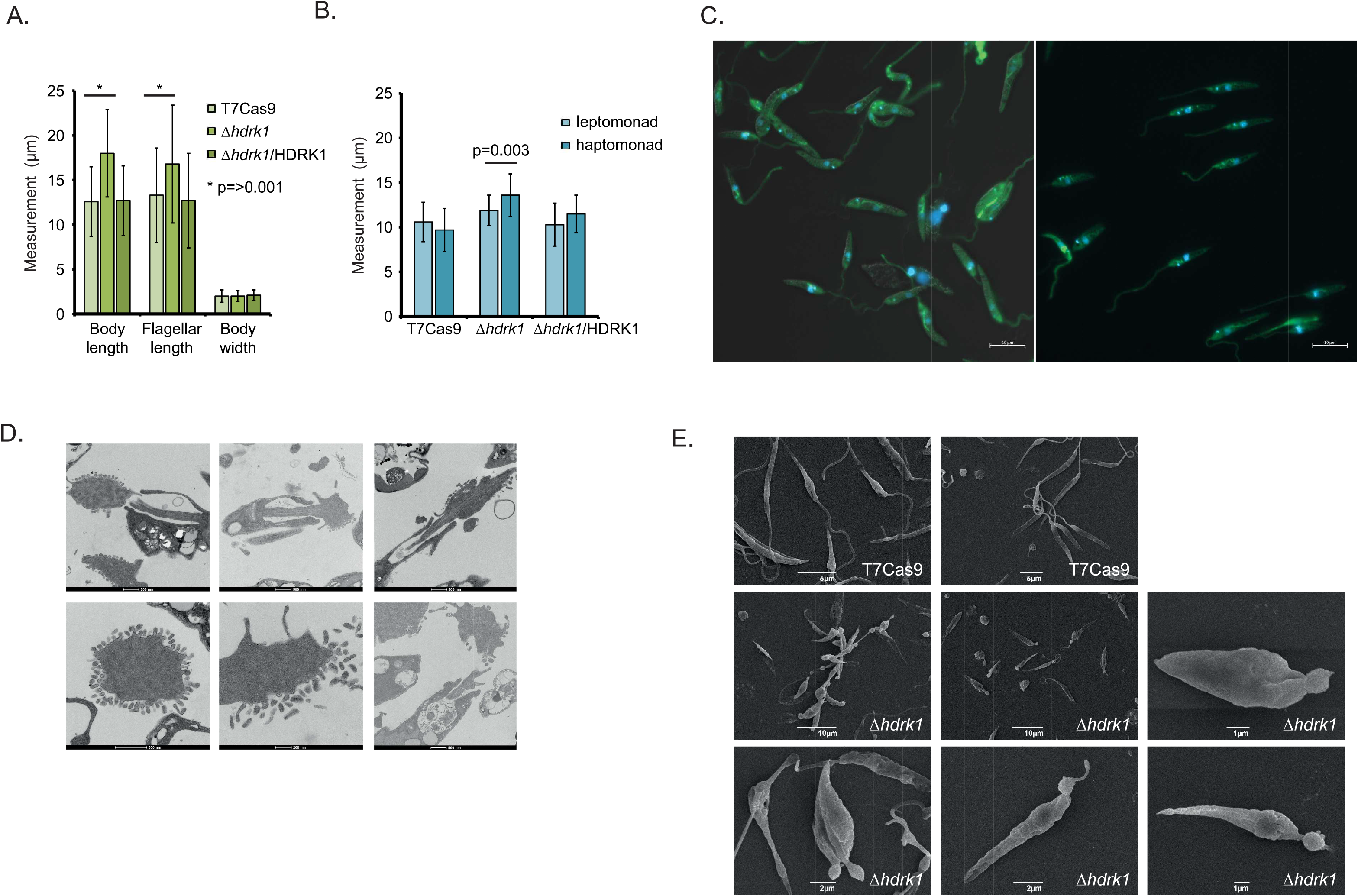
A) Bar chart showing the mean measurements, in μm, for body length, flagellar length and body width of 300 cells of mixed morphology from whole-fly dissections for each cell line. Error bars show the standard deviation. Asterisks indicate significant differences, with P < 0.001. B) Bar chart showing the mean body length measurements of leptomonad and haptomonad forms from T7Cas9, *Δhdrk1* and *Δhdrk1*/Hdrk1. Error bars show the standard deviation. n ≥ 50 for leptomonad and n > 12 for haptomonads. C) Fluorescence images of *Leishmania* expressing mNG::HDRK1, green, and stained with Hoechst 33258, blue, after 5 days at pH 5.5 in Grace’s media. Two fields of view from live-cells imaging are shown. Scale bar, 10 pm. D) TEM images *of Δhdrk1* after 7 days at low pH. The scale bar, 500 nm. E) SEM images of *Δhdrk1* after 7 days at low pH.

**Supplementary figure 5.**
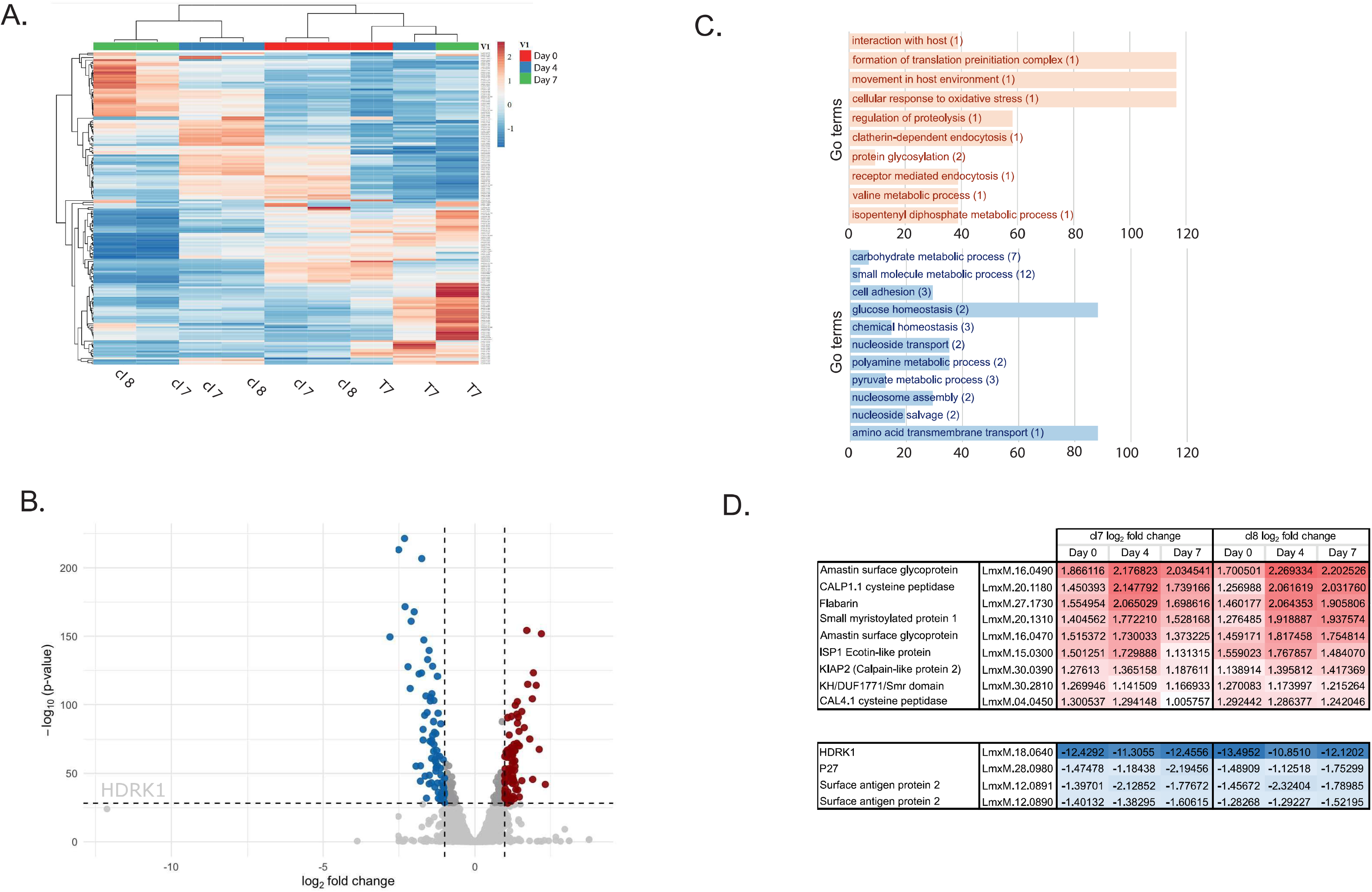
A) Heatmap of RNA-seq data collected from T7Cas9, *Δhdrk1* cl7 and *Δhdrk1* cl8 at days 0, 4 and 7 in low pH. Red represents highly expressed transcripts, and blue represents low-abundance transcripts. B) Volcano plot showing differential expression analysis comparing T7Cas9 and *Δhdrk1* cl8 at day 7 low pH. C) Slim GO term analysis for genes upregulated and downregulated in *Δhdrk1* clones at day 7 in low pH relative to T7Cas9 cells. Numbers in brackets indicate the number of genes present in the dataset with each GO term. D) Summary table indicating details of the genes that are constitutively different in *Δhdrk1* clones, irrespective of pH change, with 9 upregulated genes shown in red and 4 downregulated genes shown in blue.

**Supplementary figure 6.**
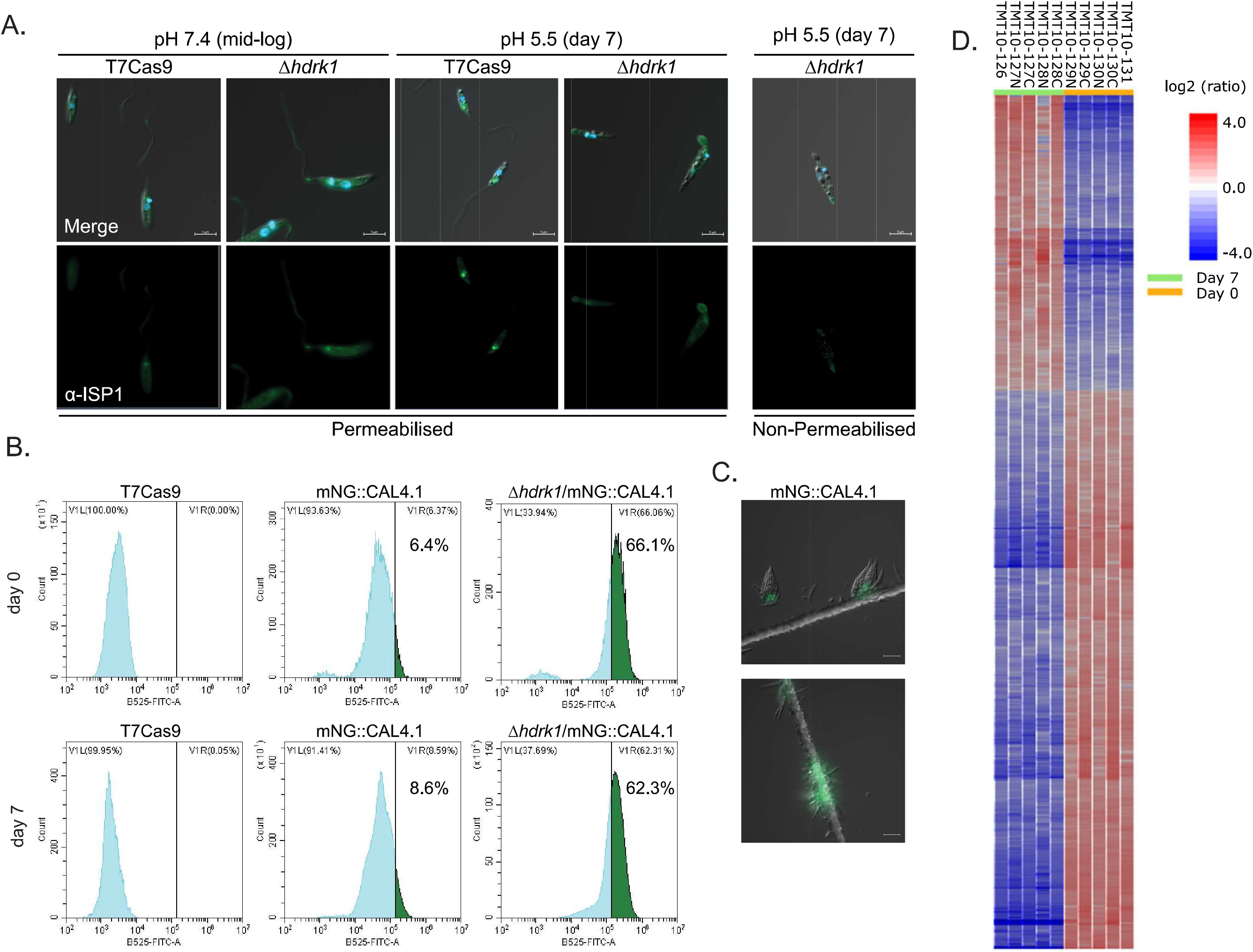
A) Immunofluorescence images showing ISP1 localisation to the bulb structure *of Δhdrk1* after 7 days at low pH, using an anti-ISP1 antibody. Hoechst is shown in blue. Scale bar, 5 µm. B) Flow cytometry of mNG::CAL4.1 expression in T7Cas9 and *Δhdrk1* at day 0 and after 7 days at low pH. C) Live-cell image of mNG::CAL4.1 in T7Cas9 cells attached to a gridded slide and covered with CyGel. Scale bar, 10 pm. D) Heatmap of whole-cell proteomic data identified usingTMT 10-plex labels, with five replicates for *Δhdrk1* day 0and five replicates for *Δhdrk1* day 7. Red indicates increased protein abundance, and blue indicates decreased abundance.

**Supplementary figure 7.**
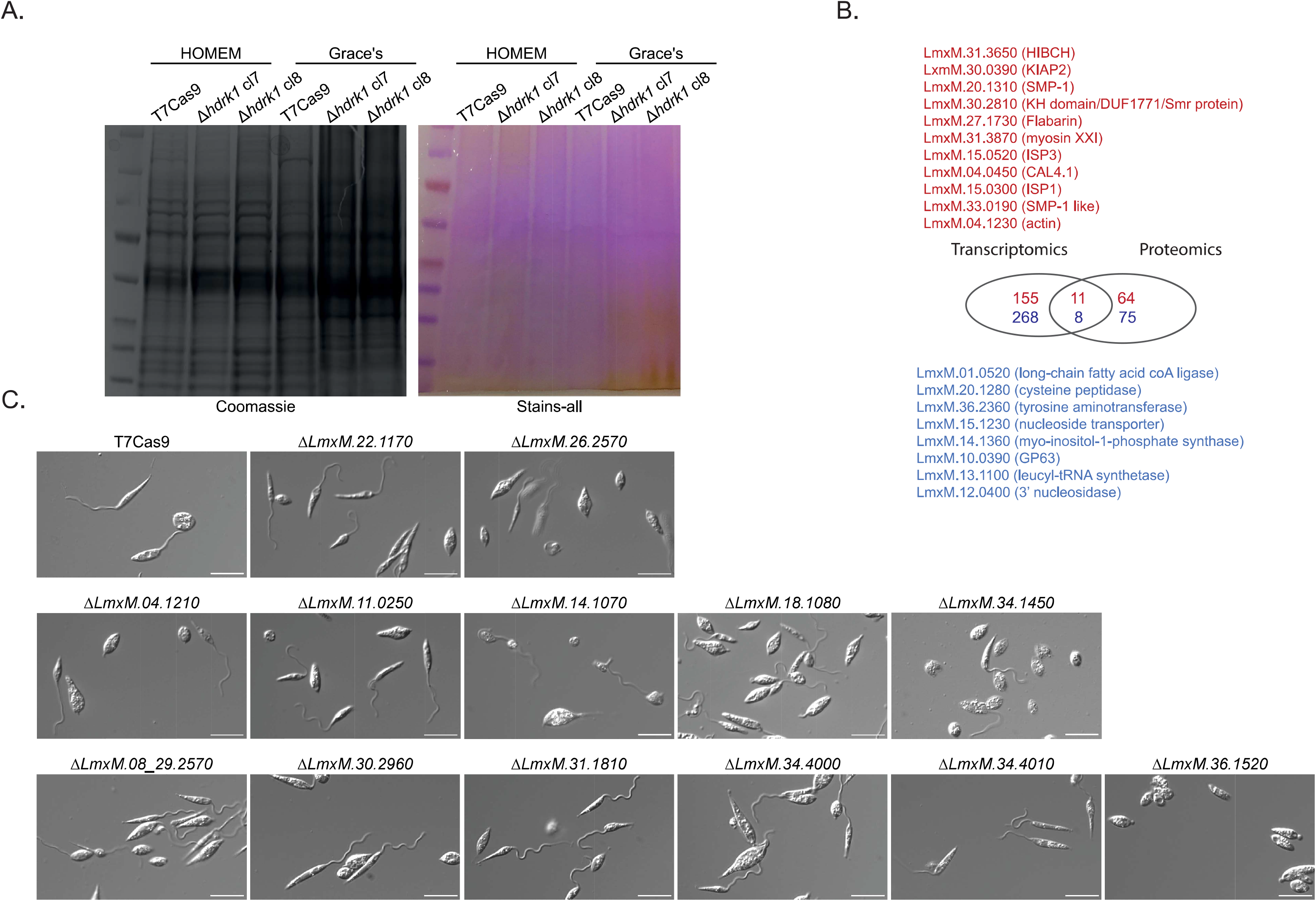
A) SDS-PAGE gel stained with Coomassie (left) and Stains-All (right). B) Overlap between transcriptomics and proteomics data. Genes upregulated in *Δhdrk1* at day 7 in low pH are shown in red text, and downregulated genes are shown in blue. C) Phasecontrast images of T7Cas9 and protein kinase mutants identified as differentially expressed by RNA-seq or differentially abundant by proteomics after 7 days in Grace’s media, pH 5.5. Kinases identified by RNA-seq were LmxM.04.1210, LmxM.11.0250, LmxM.18.1080, LmxM.22.1170, LmxM.26.2570 and LmxM.27.1780. Kinases identified by proteomics were LmxM.13.0160, LmxM.14.1070, LmxM.18.0270, LmxM.08_29.2570, LmxM.30.2960, LmxM.31.1810, LmxM.34.1050, LmxM.34.4000, LmxM.34.4010 and LmxM.36.1520. Three kinases previously found to be required for promastigote survival were not included: LmxM.27.1780, LmxM.13.0160 and LmxM.18.0270^16^. Scale bar, 10 pm.

**Supplementary figure 8.**
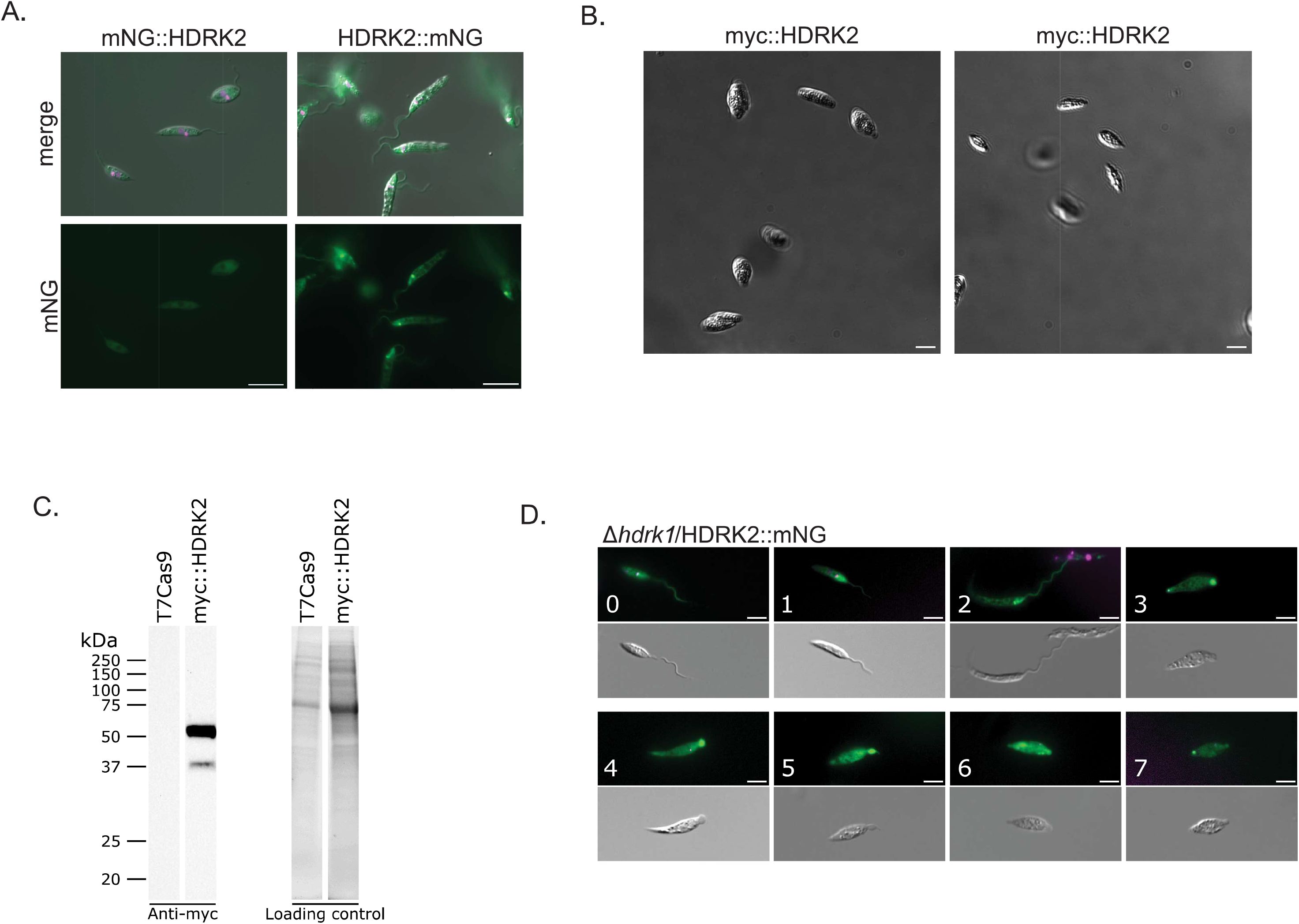
A) Live-cell images of mNG::HDRK2 and HDRK2::mNG in T7Cas9 cells after 24 h at low pH. Top, merge; bottom, mNeonGreen. Scale bar, 5 µm. B) Phase-contrast images of myc::HDRK2 after 7 days at low pH. Scale bar, 5 µm. C) Expression levels of myc::HDRK2 in M199 medium, pH 7.4. D) Live-cell images of *Δhdrk1* expressing HDRK2::mNG from day 0 to day 7 in Grace’s media. Top, mNeonGreen,green, and Hoechst,magenta, merge. Bottom: phase-contrast image. Scale bar, 5 µm.

**Supplementary figure 9.**
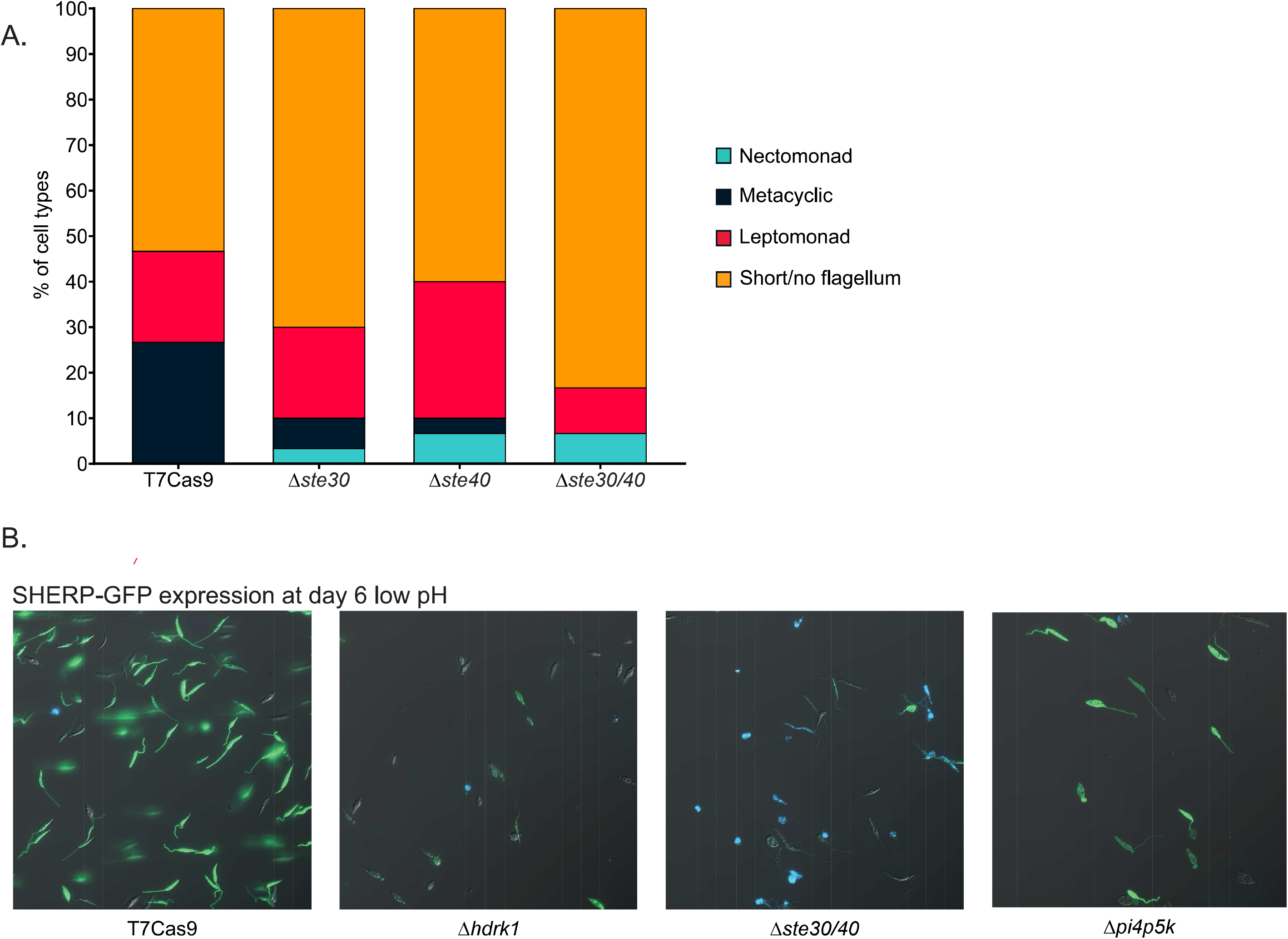
A) Bar chart showing lifecycle stages assigned by cell body width, body length and flagellum length measurements for T7Cas9, *Δste30, Δste40 and Δste30/40* after 7 days at low pH in Grace’s media. n = 30. B) Live-cell images of T7Cas9, *Δste30/40* and *Δpi4p5k* cells expressing the SHERP-GFP reporter at day 6 in Grace’s media, pH 5.5. Merged images show GFP in green and Hoechst in blue.

## Notes

### Competing Interest Statement

The authors have declared no competing interest.

## References

1. Santos, V. C., Araujo, R. N., Machado, L. A. D., Pereira, M. H. & Gontijo, N. F. The physiology of the midgut of Lutzomyia longipalpis (Lutz and Neiva 1912): pH in different physiological conditions and mechanisms involved in its control. J. Exp. Biol. 211, 2792–2798 (2008).

2. Sádlová, J. & Volf, P. Peritrophic matrix of Phlebotomus duboscqi and its kinetics during Leishmania major development. Cell Tissue Res. 337, 313–325 (2009).

3. Sacks, D. L. Metacyclogenesis in Leishmania promastigotes. Exp. Parasitol. 69, 100–103 (1989).

4. Giraud, E., Martin, O., Yakob, L. & Rogers, M. Quantifying Leishmania metacyclic promastigotes from individual sandfly bites reveals the efficiency of vector transmission. Commun. Biol. 2, 84 (2019).

5. Catta-Preta, C. M. C., Ghosh, K., Sacks, D. L. & Ferreira, T. R. Single-cell atlas of Leishmania development in sandflies reveals the heterogeneity of transmitted parasites and their role in infection. Proc. Natl. Acad. Sci. U. S. A. 121, e2406776121 (2024).

6. Gossage, S. M., Rogers, M. E. & Bates, P. A. Two separate growth phases during the development of Leishmania in sand flies: implications for understanding the life cycle. Int. J. Parasitol. 33, 1027 1034 (2003).

7. Volf, P., Hajmova, M., Sádlová, J. & Votypka, J. Blocked stomodeal valve of the insect vector: similar mechanism of transmission in two trypanosomatid models. Int. J. Parasitol. 34, 1221 1227 (2004).

8. Zilberstein, D. & Shapira, M. The role of pH and temperature in the development of Leishmania parasites. Annu. Rev. Microbiol. 48, 449–470 (1994).

9. Soête, M., Camus, D. & Dubremetz, J. F. Experimental induction of bradyzoite-specific antigen expression and cyst formation by the RH strain of Toxoplasma gondii in vitro. Exp. Parasitol. 78, 361 370 (1994).

10. Nijhout, M. M. & Carter, R. Gamete development in malaria parasites: bicarbonate-dependent stimulation by pH in vitro. Parasitology 76, 39 53 (1978).

11. Rojas, F. et al. Oligopeptide signaling through TbGPR89 drives Trypanosome quorum sensing. Cell 176, 306–317.e16 (2019).

12. Díaz, E. et al. G Protein-coupled receptors as potential intercellular communication mediators in Trypanosomatidae. Front. Cell. Infect. Microbiol. 12, 812848 (2022).

13. Parsons, M., Worthey, E. A., Ward, P. N. & Mottram, J. C. Comparative analysis of the kinomes of three pathogenic trypanosomatids: Leishmania major, Trypanosoma brucei and Trypanosoma cruzi. BMC Genomics 6, 127 (2005).

14. Jensen, B. C., Vaney, P., Flaspohler, J., Coppens, I. & Parsons, M. Unusual features and localization of the membrane kinome of Trypanosoma brucei. PLoS One 16, e0258814 (2021).

15. Cayla, M., McDonald, L., MacGregor, P. & Matthews, K. An atypical DYRK kinase connects quorum-sensing with posttranscriptional gene regulation in Trypanosoma brucei. Elife 9, e51620 (2020).

16. Baker, N. et al. Systematic functional analysis of Leishmania protein kinases identifies regulators of differentiation or survival. Nat. Commun. 12, 1244 (2021).

17. Jones, N. G. et al. Regulators of Trypanosoma brucei cell cycle progression and differentiation identified using a kinome-wide RNAi screen. PLoS Pathog. 10, e1003886 (2014).

18. Tsigankov, P. et al. Regulation dynamics of Leishmania differentiation: deconvoluting signals and identifying phosphorylation trends. Mol. Cell. Proteomics 13, 1787–1799 (2014).

19. Shaw, S. et al. Cyclic AMP signalling and glucose metabolism mediate pH taxis by African trypanosomes. Nat. Commun. 13, 603 (2022).

20. Nayak, A., Akpunarlieva, S., Barrett, M. & Burchmore, R. A defined medium for Leishmania culture allows definition of essential amino acids. Exp. Parasitol. 185, 39–52 (2018).

21. McCoy, C. J. et al. ULK4 and Fused/STK36 interact to mediate assembly of a motile flagellum. Mol. Biol. Cell 34, ar66 (2023).

22. Bates, P. A. & Tetley, L. Leishmania mexicana: induction of metacyclogenesis by cultivation of promastigotes at acidic pH. Exp. Parasitol. 76, 412–423 (1993).

23. Manning, G., Whyte, D. B., Martinez, R., Hunter, T. & Sudarsanam, S. The protein kinase complement of the human genome. Science 298, 1912–1934 (2002).

24. Crozet, P. et al. Mechanisms of regulation of SNF1/AMPK/SnRK1 protein kinases. Front. Plant Sci. 5, 190 (2014).

25. Hawley, S. A. et al. Characterization of the AMP-activated protein kinase kinase from rat liver and identification of threonine 172 as the major site at which it phosphorylates AMP-activated protein kinase. J. Biol. Chem. 271, 27879–27887 (1996).

26. Blom, N., Sicheritz-Pontén, T., Gupta, R., Gammeltoft, S. & Brunak, S. Prediction of post-translational glycosylation and phosphorylation of proteins from the amino acid sequence. Proteomics 4, 1633–1649 (2004).

27. Byrne, D. P. et al. Aurora A regulation by reversible cysteine oxidation reveals evolutionarily conserved redox control of Ser/Thr protein kinase activity. Sci. Signal. 13, eaax2713 (2020).

28. Dale, S., Wilson, W. A., Edelman, A. M. & Hardie, D. G. Similar substrate recognition motifs for mammalian AMP-activated protein kinase, higher plant HMG-CoA reductase kinase-A, yeast SNF1, and mammalian calmodulin-dependent protein kinase I. FEBS Lett. 361, 191–195 (1995).

29. Carnielli, J. B. T. et al. Chemical genetics reveals Leishmania KKT2 and CRK9 kinase activity is required for cell cycle progression. PLoS Pathog. 22, e1014194 (2026).

30. McNiven, C., Carnielli Trindade, J. B. T., Geoghegan, V., Faria, J. R. C. & Mottram, J. C. CRISPR-Cas9 precision editing of kinetochore protein phosphosite codons in Leishmania mexicana. Front. Cell. Infect. Microbiol. 16, 1788564 (2026).

31. Walters, L. L. Leishmania differentiation in natural and unnatural sand fly hosts 1. J. Eukaryot. Microbiol. 40, 196–206 (1993).

32. Yanase, R. et al. Discovery of essential kinetoplastid-insect adhesion proteins and their function in Leishmania-sand fly interactions. Nat. Commun. 15, 6960 (2024).

33. Morrison, L. S. et al. Ecotin-like serine peptidase inhibitor ISP1 of Leishmania major plays a role in flagellar pocket dynamics and promastigote differentiation. Cell. Microbiol. 14, 1271–1286 (2012).

34. Di-Blasi, T. et al. The flagellar protein FLAG1/SMP1 is a candidate for Leishmania-sand fly interaction. Vector Borne Zoonotic Dis. 15, 202–209 (2015).

35. Giordana, L. et al. Molecular and functional characterization of two malic enzymes from Leishmania parasites. Mol. Biochem. Parasitol. 219, 67–76 (2018).

36. Fochler, S., Walker, B. J., Wheeler, R. J. & Gluenz, E. Divergent protein kinase A contributes to the regulation of flagellar waveforms in Leishmania mexicana. J. Cell Sci. 139, (2026).

37. Hassan, P., Fergusson, D., Grant, K. M. & Mottram, J. C. The CRK3 protein kinase is essential for cell cycle progression of Leishmania mexicana. Mol. Biochem. Parasitol. 113, 189–198 (2001).

38. Owino, B. O. et al. Discovery of a novel flagellar filament system underpinning Leishmania adhesion to surfaces. Curr. Biol. 35, 2837–2847.e4 (2025).

39. Lemmon, M. A. & Ferguson, K. M. Signal-dependent membrane targeting by pleckstrin homology (PH) domains. Biochem. J. 350, 1–18 (2000).

40. Abramson, J. et al. Accurate structure prediction of biomolecular interactions with AlphaFold 3. Nature 630, 493–500 (2024).

41. van Kempen, M. et al. Fast and accurate protein structure search with Foldseek. Nat. Biotechnol. 42, 243–246 (2024).

42. Gumerov, V. M. et al. Amino acid sensor conserved from bacteria to humans. Proc. Natl. Acad. Sci. U. S. A. 119, e2110415119 (2022).

43. Tohidifar, P. et al. The unconventional cytoplasmic sensing mechanism for ethanol chemotaxis in Bacillus subtilis. mBio 11, e02177–20 (2020).

44. Sádlová, J. et al. The stage-regulated HASPB and SHERP proteins are essential for differentiation of the protozoan parasite Leishmania major in its sand fly vector, Phlebotomus papatasi. Cell. Microbiol. 12, 1765–1779 (2010).

45. Knuepfer, E., Stierhof, Y. D., McKean, P. G. & Smith, D. F. Characterization of a differentially expressed protein that shows an unusual localization to intracellular membranes in Leishmania major. Biochem. J. 356, 335–344 (2001).

46. Denecke, S. et al. Adhesion of Crithidia fasciculata promotes a rapid change in developmental fate driven by cAMP signaling. mSphere 9, e0061724 (2024).

47. Skalický, T. et al. Extensive flagellar remodeling during the complex life cycle of Paratrypanosoma, an early-branching trypanosomatid. Proc. Natl. Acad. Sci. U. S. A. 114, 11757–11762 (2017).

48. Filosa, J. N. et al. Dramatic changes in gene expression in different forms of Crithidia fasciculata reveal potential mechanisms for insect-specific adhesion in kinetoplastid parasites. PLoS Negl. Trop. Dis. 13, e0007570 (2019).

49. Owino, B. O. et al. Identification of a conserved gene family with an essential role in Leishmania parasite-insect vector adhesion. Proc. Natl. Acad. Sci. U. S. A. 123, e2603653123 (2026).

50. Tull, D. et al. SMP-1, a member of a new family of small myristoylated proteins in kinetoplastid parasites, is targeted to the flagellum membrane in Leishmania. Mol. Biol. Cell 15, 4775–4786 (2004).

51. Halliday, C. et al. Role for the flagellum attachment zone in Leishmania anterior cell tip morphogenesis. PLoS Pathog. 16, e1008494 (2020).

52. Saldivia, M., Ceballos-Pérez, G., Bart, J.-M & Navarro, M. The AMPKal pathway positively regulates the developmental transition from proliferation to quiescence in Trypanosoma brucei. Cell Rep. 17, 660–670 (2016).

53. Nolan, D. P., Rolin, S., Rodriguez, J. R., Van Den Abbeele, J. & Pays, E. Slender and stumpy bloodstream forms of Trypanosoma brucei display a differential response to extracellular acidic and proteolytic stress. Eur. J. Biochem. 267, 18–27 (2000).

54. Shoeran, G., Anand, N., Kaur, U., Goyal, K. & Sehgal, R. Identification and characterization of yeast SNF1 kinase homologs in Leishmania major. Front. Mol. Biosci. 12, 1567703 (2025).

55. Myburgh, E. et al. TORC1 is an essential regulator of nutrient-controlled proliferation and differentiation in Leishmania. EMBO Rep. 25, 1075–1105 (2024).

56. Glaser, T. A., Baatz, J. E., Kreishman, G. P. & Mukkada, A. J. pH homeostasis in Leishmania donovani amastigotes and promastigotes. Proc. Natl. Acad. Sci. U. S. A. 85, 7602–7606 (1988).

57. Yang, X. et al. The α subunit of AMP-activated protein kinase is critical for the metabolic success and tachyzoite proliferation of Toxoplasma gondii. Microb. Biotechnol. 17, e14455 (2024).

58. Beneke, T. et al. A CRISPR Cas9 high-throughput genome editing toolkit for kinetoplastids. R Soc Open Sci 4, 170095 (2017).

59. Spence, K. Investigating the role of the SNARE protein, Tlg2, in Leishmania mexicana. (University of York, 2022).

60. Volf, P. & Volfova, V. Establishment and maintenance of sand fly colonies. J. Vector Ecol. 36 Suppl 1, S1–9 (2011).

61. Myskova, J., Votypka, J. & Volf, P. Leishmania in sand flies: comparison of quantitative polymerase chain reaction with other techniques to determine the intensity of infection. J. Med. Entomol. 45, 133–138 (2008).

62. Yanase, R. et al. Formation and three-dimensional architecture of Leishmania adhesion in the sand fly vector. Elife 12, e84552 (2023).

63. Grewal, J. S., Catta-Preta, C. M. C., Brown, E., Anand, J. & Mottram, J. C. Evaluation of clan CD C11 peptidase PNT1 and other Leishmania mexicana cysteine peptidases as potential drug targets. Biochimie 166, 150–160 (2019).

64. Andrews, S. (2010) FastQC: A quality control tool for high throughput sequence data. https://www.bioinformatics.babraham.ac.uk/projects/fastqc/.

65. Ewels, P., Magnusson, M., Lundin, S. & Käller, M. MultiQC: summarize analysis results for multiple tools and samples in a single report. Bioinformatics 32, 3047–3048 (2016).

66. Kim, D., Paggi, J. M., Park, C., Bennett, C. & Salzberg, S. L. Graph-based genome alignment and genotyping with HISAT2 and HISAT-genotype. Nat. Biotechnol. 37, 907–915 (2019).

67. Love, M. I., Huber, W. & Anders, S. Moderated estimation of fold change and dispersion for RNA-seq data with DESeq2. Genome Biol. 15, 550 (2014).

68. Zhu, A., Ibrahim, J. G. & Love, M. I. Heavy-tailed prior distributions for sequence count data: removing the noise and preserving large differences. Bioinformatics 35, 2084–2092 (2019).

69. Metsalu, T. & Vilo, J. ClustVis: a web tool for visualizing clustering of multivariate data using Principal Component Analysis and heatmap. Nucleic Acids Res. 43, W566–70 (2015).

70. Corpet, F. Multiple sequence alignment with hierarchical clustering. Nucleic Acids Res. 16, 10881–10890 (1988).

