## Supplementary information for "A pH-dependent protein kinase cascade regulates divergent differentiation of *Leishmania* in the sand fly"

**Cell Lines:** List of cell lines used in this study including unique identifiers and resistance markers present.

| **Cell line identifier** | **Description** | **Gene of interest** | **Resistance Markers** | **Parental line** |
| --- | --- | --- | --- | --- |
| JM7607 | *Δhdrk1* cl7 | LmxM.18.0640 | PURO/PURO | T7Cas9 |
| JM7608 | *Δhdrk1* cl8 | LmxM.18.0640 | PURO/PURO | T7Cas9 |
| JM7800 | *Δhdrk1::*HDRK1 SSU *addback A3* | LmxM.18.0640 | PURO/PURO/G418 | JM7608 |
| JM7672 | *Δhdrk1* + SHERP-GFP reporter | - | PURO/PURO/G418 | JM7608 |
| JM7669 | T7cas9 + SHERP-GFP reporter | - | PURO/PURO/G418 | T7Cas9 |
| JM8914 | K38a CL1 precision edit of LmxM.18.0640 (C11) | LmxM.18.0640 | - | T7Cas9 |
| JM8913 | K38 Sil CL7 precision edit of LmxM.18.0640 (C11) | LmxM.18.0640 | - | T7Cas9 |
| ST2 | HDRK1 (LmxM.18.0640) precision edit D132A, D150A CL2 | LmxM.18.0640 | - | T7Cas9 |
| ST3 | HDRK1 (LmxM.18.0640) precision edit D132A, D150A CL3 | LmxM.18.0640 | - | T7Cas9 |
| JM8911 | LmxM.18.0640 (HDRK1) precision edit D132Sil, D150Sil Clone DD2 | LmxM.18.0640 | - | T7Cas9 |
| JM8880 | LmxM.18.0640 HDRK1 C166A Catalytic Mutant homozygous Clone 8 | LmxM.18.0640 | - | T7Cas9 |
| JM8879 | LmxM.18.0640 HDRK1 C166A Catalytic Mutant homozygous Clone 6 | LmxM.18.0640 | - | T7Cas9 |
| JM8881 | LmxM.18.0640 HDRK1 C166 Silent mutant homozygous Clone 3 | LmxM.18.0640 | - | T7Cas9 |
| St71 | HDRK1 (LmxM.18.0640) precision edit T164T (Sil) clone 1.1 | LmxM.18.0640 | - | T7Cas9 |
| St72 | HDRK1 (LmxM.18.0640) precision edit T164T (Sil) clone 1.4 | LmxM.18.0640 | - | T7Cas9 |
| St74 | HDRK1 (LmxM.18.0640) precision edit T164A clone 2.1 | LmxM.18.0640 | - | T7Cas9 |
| St80 | HDRK1 (LmxM.18.0640) precision edit T164A clone 2.9 | LmxM.18.0640 | - | T7Cas9 |
| St76 | HDRK1 (LmxM.18.0640) precision edit T164D clone 3.4 | LmxM.18.0640 | - | T7Cas9 |
| St75 | HDRK1 (LmxM.18.0640) precision edit T164D clone 3.1 | LmxM.18.0640 | - | T7Cas9 |
| JM8633 | mNeon::HDRK1 (N term) | LmxM.18.0640 | BLA | T7Cas9 |
| JM8634 | HDRK1::mNeon (C term) | LmxM.18.0640 | BLA | T7Cas9 |
| ST35 | myc::HDRK1 (N-terminal) | LmxM.18.0640 | PURO/BLA | T7Cas9 |
| JM7616 | HDRK1 null (cl8) + mNeon::ISP1 | LmxM.15.0300 | BLA | JM7608 |
| JM7617 | HDRK1 null (cl8) + ISP1 ::mNeon | LmxM.15.0300 | BLA | JM7608 |
| JM7649 | mNeon::ISP1 | LmxM.15.0300 | BLA | T7Cas9 |
| JM8295 | *Δhdrk1* CL 8 mNeon::CAL4.1 (n term) | LmxM.04.0450 | PURO/PURO BLA/G418 | JM7608 |
| JM8296 | mNeon::CAL4.1 (n term) | LmxM.04.0450 | BLA/G418 | T7Cas9 |
| JM8554 | *Δhdrk2 NEK17 KO (LmxM.36.1520) (BLA ONLY) Clone NA2* | LmxM.36.1520 | BLA | T7Cas9 |
| JM8555 | *Δhdrk2 NEK17 KO (LmxM.36.1520) (BLA ONLY) Clone NB1* | LmxM.36.1520 | BLA | T7Cas9 |
| JM8635 | mNeon::HDRK2 (N term) | LmxM.36.1520 | Puro | T7Cas9 |
| JM8636 | HDRK2::mNeon (C term) | LmxM.36.1520 | Puro | T7Cas9 |
| JM8637 | *Δhdrk2 + mNeon HDRK1 NB1 pop* | LmxM.36.1520 | Bla/Bla Puro | JM8555 |
| JM8638 | *Δhdrk2 + mNeon HDRK1 NB2 pop* | LmxM.36.1520 | Bla/Bla Puro | JM8556 |
| JM8639 | Δ*ste30/40* (LmxM.30.1840/1830) mNeon::HDRK1 | LmxM.18.0640 | BLA/BLA/G418 | JM8550 |
| JM8550 | *Δste30/40 SB2 clone* | LmxM.30.1830/40 | BLA/BLA | T7Cas9 |
| JM8551 | *Δ*ste30/40 *SA5 clone* | LmxM.30.1830/40 | BLA/BLA | T7Cas9 |
| ST37 | *STE 30/40/08.0600 Triple null* | LmxM.30.1830/40, LmxM.08.0660 | bla/bla/puro/g418 | JM8550 |
| ST30 | *Δ*ste30/40 *SB2 clone (JM8550 - BLA) + myc::HDRK1 puro neo* | LmxM.30.1830/40 | bla/bla/puro/g418 | JM8550 |

**Primers:** Sequences and unique identifiers for each of the oligonucleotides used in this study

| **Edit** | **Gene taget** | **Oligo number** | **Oligo description** | **Oligo sequence** |
| --- | --- | --- | --- | --- |
| HDRK1 Null mutant | LmxM.18.0640 | OL7588 | Upstream Forward | AACGTCGTCGAGGACGGTCATCACAGCCACgtataatgcagacctgctgc |
|  |  | OL9080 | Downstream Reverse | CTTTCTACGGAGCTTCACTCCCCCGCCCCGccaatttgagagacctgtgc |
|  |  | OL7572 | 5' guide | gaaattaatacgactcactataggATACACGAACAAGTAGCTACgttttagagctagaaatagc |
|  |  | OL9109 | 3'guide | gaaattaatacgactcactataggTGGCAGATGATGGGGGAGATgttttagagctagaaatagc |
|  |  | OL9671 | Internal ORF Fw 490bp | GGAGAACATGGAGGACCAGA |
|  |  | OL9700 | Internal ORF Rev 490bp | GTTTCGATCCTCGAACGGTA |
|  |  | OL9369 | Integration Rev (in 5' UTR of all repair cassettes) | GCAGCAGGTCTGCATTATAC |
|  |  | OL9729 | Integration fwd (use with OL9369) | CGGCGACAGACACCTATTCT |
| HDRK1 Addback | LmxM.18.0640 | OL 11563 | 856bp amplification with overlaps for gibson assembly into pRib vector | ttgagccgtccaccgtagccTCGAGATGGATCCATCTGTCATG |
|  |  | OL11564 | 856bp amplification with overlaps for gibson assembly into pRib vector | tcttgcttaaggcgcgccgcGGCCGCTTACGTGGAGAGTAACCTC |
|  |  | OL11543 | HDRK1 addback colony PCR F (Should amplify around 470bp from pgl2398 plus insert) | ATGAGGCAGCGGAAAGGAAT |
|  |  | OL11544 | HDRK1 addback colony PCR R | AAACACGGCTAAGCGAAAGC |
|  |  | OL11545 | HDRK1 addback sequencing F (Should amplify around 274bp from pgl2398 plus insert) | TCGAGTAACAATTGCCCGCT |
|  |  | OL11546 | HDRK1 addback sequencing R | TGCGAATTTGCAACCTGAGC |
| HDRK1 N terminal tag | LmxM.18.0640 | OL7604 | Upstream reverse | GTACTGGCCCACCATGACAGATGGATCCATactacccgatcctgatccag |
| HDRK1 C terminal tag | LmxM.18.0640 | OL12352 | Downstream forward primer | ATTGAGTGGAATCCGAGGTTACTCTCCACGggttctggtagtggttccgg |
| HDRK1 Myc only N term tag | LmxM.18.0640 | OL11953 | Rev to amplify only resistance marker and 3x myc (No mNeon) | GTACTGGCCCACCATGACAGATGGATCCATagaaccggaaccggaaccac |
| SHERP-GFP reporter | SHERP | OL11254 | pTNEO For addition of G418 to pSSU GFP 3'UTR sherp plasmid (waldrad lab) | taatgtgtacgtatatgacaatgggatcggccattgaac |
|  |  | OL11255 | pTNEO For addition of G418 to pSSU GFP 3'UTR sherp plasmid (waldrad lab) | acacacgcacacaagggcatgaagaactcgtcaagaaggc |
|  |  | OL11256 | pSSU GFP 3' UTR Sherp g418 For sequencing from sherp plasmid into G418 (92bp) | GTGGGTGTTTCCGGTGATGT |
|  |  | OL11257 | pSSU GFP 3' UTR Sherp g418 For sequencing from sherp plasmid into G418 (92bp) | GAACCTGCGTGCAATCCATC |
|  |  | OL11258 | pSSU GFP 3' UTR Sherp g418 For sequencing from G418 into sherp plasmid (192) | GCTTCCTCGTGCTTTACGGTATC |
|  |  | OL11259 | pSSU GFP 3' UTR Sherp g418 For sequencing from G418 into sherp plasmid (192) | CCTCTGCCTCACATCTCGCA |
| HDRK1 K38A precision edit | LmxM.18.0640 | OL14403 | DS precision edit repair F | GATCCATCTGTCATGGTGGGCCAGTACACGCTGGGGAAGCAGATCGGCTCCGGCAACTTCTCCAAGGTGCGtCTcGGtACaGAcCCcCAgGGtCGCACATGGGCCATCAA |
|  |  | OL14404 | DS precision edit repair R | TTCTGCTGACGCAGACTGCGCATGATGGCGACTTCACGGAGCATCTGGTCCTCCATGTTCTCtTTCTTaAGcCGcCGtTTgTCCACGATtTTGATGGCCCATGTGCGaCC |
|  |  | OL14405 | DS precision edit repair F | GATCCATCTGTCATGGTGGGCCAGTACACGCTGGGGAAGCAGATCGGCTCCGGCAACTTCTCCAAGGTGCGtCTcGGtACaGAcCCcCAgGGtCGCACATGGGCCATCGC |
|  |  | OL14406 | DS precision edit repair R | TTCTGCTGACGCAGACTGCGCATGATGGCGACTTCACGGAGCATCTGGTCCTCCATGTTCTCtTTCTTaAGcCGcCGtTTgTCCACGATGGCGATGGCCCATGTGCGaCC |
|  |  | OL14407 | Precision edit 5' g | GAAATTAATACGACTCACTATAGGCGACTGGGCACCGATCCGCAGTTTTAGAGCTAGAAATAGC |
|  |  | OL14408 | Precision edit 3' guide g | GAAATTAATACGACTCACTATAGGGGATAAGCGCCGCCTAAAGAGTTTTAGAGCTAGAAATAGC |
|  |  | OL14409 | Edit sprific primer 682bp F | TCTCGGTACAGACCCCCAG |
|  |  | OL14410 | Edit sprific primer 682bp R | CTCCATTGAGATGCGCTTGC |
|  |  | OL14411 | 903bp region (compatible for sequencing or Bsr1 digestion diagnostic) F | TTCCTCTCCCCCTCTTCACA |
|  |  | OL14412 | 903bp region (compatible for sequencing or Bsr1 digestion diagnostic) R | GGAGAGTAACCTCGGATTCCA |
| HDRK1 D132A,D150A presicion edit | LmxM.18.0640 | OL14413 | LmxM.18.0640 (D132&D15 sil) DS precision edit repair | CGCTTCGACGAGCCGACGGCGCGTCGGTACTTCCACCAGCTCATCGCAGGCATGTAtTAtTGtCAtTCcAAaGGtTTCGCCCACCGCGATCTGAAGCCGGAGAACCTGTTGCTCGACGCG |
|  |  | OL14414 | LmxM.18.0640 (D132&D15 sil) DS precision edit repair | GCACCTCCGGTGCAACGTAGTTTGGCGTGCCGCAGATGGTCTGAAGCAGCACATCtTGtTGgAGaTTaCTaAGaCCAAAATCGGAAATCTTCAGTGTGCCGTTCGCGTCGAGCAACAGGT |
|  |  | OL14415 | LmxM.18.0640 (D132A&D150A) DS precision edit repair | CGCTTCGACGAGCCGACGGCGCGTCGGTACTTCCACCAGCTCATCGCAGGCATGTAtTAtTGtCAtTCcAAaGGtTTCGCCCACCGCGCCCTGAAGCCGGAGAACCTGTTGCTCGACGCG |
|  |  | OL14416 | LmxM.18.0640 (D132A&D150A) DS precision edit repair | GCACCTCCGGTGCAACGTAGTTTGGCGTGCCGCAGATGGTCTGAAGCAGCACATCtTGtTGgAGaTTaCTaAGaCCAAAGGCGGAAATCTTCAGTGTGCCGTTCGCGTCGAGCAACAGGT |
|  |  | OL14417 | LmxM.18.0640 (D132A&D150A) Precision edit 5' | GAAATTAATACGACTCACTATAGGATGTACTACTGCCACTCGAAGTTTTAGAGCTAGAAATAGC |
|  |  | OL14418 | LmxM.18.0640 (D132A&D150A) Precision edit 3' guide | GAAATTAATACGACTCACTATAGGTGGGCTCAGCAACCTTCAGCGTTTTAGAGCTAGAAATAGC |
|  |  | OL14419 | LmxM.18.0640 (D132A&D150A) Edit specific primer 455bp | TGTCATTCCAAAGGTTTCGCC |
|  |  | OL14420 | LmxM.18.0640 (D132A&D150A) Edit sprific primer 455bp | GCCGCTTACGTGGAGAGTAA |
| HDRK1 C166A precision edit | LmxM.18.0640 | OL14340 | LmxM.18.0640 (C11/HDRK1) C166A repair primer F | GCTCGACGCGAACGGCACACTGAAGATTTCCGACTTTGGGCTCAGCAACCTTCAGCAGGATGTCTTGCTACAAACAATTGCAGGTACCCCGAATTATGTAGC |
|  |  | OL14341 | LmxM.18.0640 (C11/HDRK1) C166 silent repair primer F | GCTCGACGCGAACGGCACACTGAAGATTTCCGACTTTGGGCTCAGCAACCTTCAGCAGGATGTCTTGCTACAAACAATTTGTGGTACCCCGAATTATGTAGC |
|  |  | OL14342 | LmxM.18.0640 (C11/HDRK1) C166 repair primer R | GCAGCTCCAGATGTCCGCCGAGAGTCCGTTGTAGCCGCGCTCCATGAGCACCTCTGGAGCTACATAATTCGGGGTACC |
|  |  | OL14345 | LmxM.18.0640 (C11/HDRK1) C166 guide 1 | gaaattaatacgactcactataggGTGCTGCTTCAGACCATCTGgttttagagctagaaatagc |
|  |  | OL14346 | LmxM.18.0640 (C11/HDRK1) C166 guide 2 | gaaattaatacgactcactataggCACGCCAAACTACGTTGCACgttttagagctagaaatagc |
| HDRK1 T164 precision edit | LmxM.18.0640 | Pr49 | 5' guide | gaaattaatacgactcactataggTGGGCTCAGCAACCTTCAGCgttttagagctagaaatagc |
|  |  | Pr50 | 3' guide | gaaattaatacgactcactataggCACGCCAAACTACGTTGCACgttttagagctagaaatagc |
|  |  | Pr51 | T164T Fw (Sil) | AAGCCGGAGAACCTGTTGCTCGACGCGAACGGCACACTGAAGATTTCCGACTTTGGcCTgAGtAAtCTcCAaCAaGATGTGCTGCTTCAGACgATCTGCG |
|  |  | Pr52 | T164T Rv | CTCCAGATGTCCGCCGAGAGTCCGTTGTAGCCGCGCTCCATGAGCACCTCTGGCGCGACATAATTCGGGGTGCCGCAGATCGTCTGAAGCAGC |
|  |  | Pr53 | T164A Fw (inactive) | AAGCCGGAGAACCTGTTGCTCGACGCGAACGGCACACTGAAGATTTCCGACTTTGGcCTgAGtAAtCTcCAaCAaGATGTGCTGCTTCAGgccATCTGCG |
|  |  | Pr54 | T164A Rev (inactive) | CTCCAGATGTCCGCCGAGAGTCCGTTGTAGCCGCGCTCCATGAGCACCTCTGGCGCGACATAATTCGGGGTGCCGCAGATggcCTGAAGCAGC |
|  |  | Pr55 | T164D F (active) | AAGCCGGAGAACCTGTTGCTCGACGCGAACGGCACACTGAAGATTTCCGACTTTGGcCTgAGtAAtCTcCAaCAaGATGTGCTGCTTCAGgacATCTGCG |
|  |  | Pr56 | T164D R | CTCCAGATGTCCGCCGAGAGTCCGTTGTAGCCGCGCTCCATGAGCACCTCTGGCGCGACATAATTCGGGGTGCCGCAGATgtcCTGAAGCAGC |
|  |  | Pr57 | 415bp | ACTTCTCCAAGGTGCGACTG |
|  |  | Pr58 | 415bp | TGTTGGAGATTACTCAGGCCA |
| ISP1 N/C terminal tag | LmxM.15.0300 | OL11263 | ISP1 (LmxM.15.0300) Upstream forward primer | CACCGGCACCTCTCCTCCCTCCCCCCTCCCgtataatgcagacctgctgc |
|  |  | OL11264 | ISP1 (LmxM.15.0300) Upstream reverse primer | GTACGGGGCCTCGATCTTGCAGTACGACATactacccgatcctgatccag |
|  |  | OL11265 | ISP1 (LmxM.15.0300) 5' guide | gaaattaatacgactcactataggGTGTGCTTCAGACGTAAGCTgttttagagctagaaatagc |
|  |  | OL11266 | ISP1 (LmxM.15.0300) Downstream forward | GGAGGCCGGCAGATGCAGGCAGCCACAGAGggttctggtagtggttccgg |
|  |  | OL11267 | ISP1 (LmxM.15.0300) Downstream reverse | GCGGGCGACCGCTAAGGTGAGAAAGCACAAccaatttgagagacctgtgc |
|  |  | OL11268 | ISP1 (LmxM.15.0300) 3' guide | gaaattaatacgactcactataggCGAGCGGGCGAGCTCGAAATgttttagagctagaaatagc |
| Cysteiene peptidase N term tag | LmxM.27.500 | OL13180 | Forward no barcode | GCCCGAACACGAATGGTGGGGACACGTCCCgtataatgcagacctgctgc |
|  |  | OL13181 | Upstream Reverse | ATGTCTTCGTCGGGTTGAAAAAATGAACATactacccgatcctgatccag |
|  |  | OL13178 | Upstream sgRNA primer | gaaattaatacgactcactataggGTTCTGGCGCCACGATACGAgttttagagctagaaatagc |
| CAL4.1 N term tag | LmxM.04.0450 | OL7033 | Upstream Forward | TAATACGACTCACTATAAAACTGGAAGTTGTATAGCCGGTCCTCTCCACCCGCCCACCCACCCACCCCAgtataatgcagacctgctgc |
|  |  | OL7034 | upstream Reverse | CGGCTTGTCGAGAGGGGGGAGGCACCCCCCccaatttgagagacctgtgc |
|  |  | OL7035 | 5' guide | gaaattaatacgactcactataggATGTGCAGGAGGTGTGTGTAgttttagagctagaaatagc |
| HDRK2 / NEK17 Tagging/knockout | LmxM.36.1520 | OL13200 | Non Barcoded Upsteam forward | TTCTGGTGTGTTGAGCACCTATCAGCGCACgtataatgcagacctgctgc |
|  |  | OL13201 | Upstream Reverse | CATGGCGTCGCCGGCGTTACCCCCCGACATactacccgatcctgatccag |
|  |  | OL13202 | upsteam guide | gaaattaatacgactcactataggTGAACGAGGGAAAGTGTATAgttttagagctagaaatagc |
|  |  | OL14035 | Downstream forward primer | GACGACATGTCTCCTCTGCCCGCAAAGGAGggttctggtagtggttccgg |
|  |  | OL14036 | Downstream reverse primer | TGCCTTCCGCTTCTTGTGTTGTCACATCCTccaatttgagagacctgtgc |
|  |  | OL14037 | 3' sgRNA primer | gaaattaatacgactcactataggTCTACAACGCGCAGAAGAAAgttttagagctagaaatagc |
|  |  | OL9189 | Internal CDS Fw | CTCTGCTTCTTCACGGTTCC |
|  |  | OL9210 | Internal CDS Rv | AGGCAGGTCTGCTGTTCCTA |
|  |  | OL9766 | Integration Fwd (use with OL9369) | GACGCTTTTCTTGCTGCTCT |
| STE 30/40 Null | LmxM.30.1830/40 | OL11984 | LmxM.30.1840 Forward | CAAGGCAGGTAAGTGACCCCCCCCCCTCCCgtataatgcagacctgctgc |
|  |  | OL11987 | LmxM.30.1840 guide | gaaattaatacgactcactataggGGAATAGCTGCCGATGCCCTgttttagagctagaaatagc |
|  |  | OL6180 | LmxM.30.1830 Downstream Rev | AGCTGTTCCAGTGGCATTCTCCGCTCGGCTccaatttgagagacctgtgc |
|  |  | OL6181 | LmxM.30.1830 3' guide | gaaattaatacgactcactataggAATGGCCTAGCCGGCAAAACgttttagagctagaaatagc |
|  |  | OL14009 | Forward outside ORF | CGTCAGGTGCACATACCACA |
|  |  | OL14010 | Reverse outside ORF | GAGACACCGACATCCAGAGG |
| HDRK1 protein expression | LmxM.18.0640 | OL12207 | AML0218 pUC-sp LmxM.18.0640 | TCCAGGGACCAGCAATGGACCCGTCTGTAATGGTTG |
|  |  | OL12208 | AML0218 pUC-sp LmxM.18.0640 | TGAGGAGAAGGCGCGTTACGTGGACAACAGACGCG |

**Phenotypes linked to kinase mutants at day 7 low pH (figure 1d):**

| Counts | T7Cas9 | Δ*hdrk1* | Δ*25.1560* | Δ*21.0823* | Δ*pi3k* | Δ*rdk1* | Δ*19.1470* | Δ*30.1830* | Δ*pi4p5k* | Δ*30.1840* | Δ*07.0250* |
| --- | --- | --- | --- | --- | --- | --- | --- | --- | --- | --- | --- |
| procyclic promastigote | 5 | 37 | 3 | 2 | 5 | 1 | 7 | 6 | 3 | 4 | 13 |
| elongated nectomonad | 7 | 10 | 9 | 9 | 24 | 36 | 6 | 9 | 29 | 22 | 3 |
| short nectomonads | 35 | 3 | 36 | 34 | 17 | 9 | 31 | 33 | 17 | 21 | 33 |
| metacyclic promastigote | 3 | 0 | 2 | 5 | 4 | 4 | 6 | 2 | 1 | 3 | 1 |
| % | T7Cas9 | Δ*hdrk1* | Δ*25.1560* | Δ*21.0823* | Δ*pi3k* | Δ*rdk1* | Δ*19.1470* | Δ*30.1830* | Δ*pi4p5k* | Δ*30.1840* | Δ*07.0250* |
| procyclic promastigote | 10 | 74 | 6 | 4 | 10 | 2 | 14 | 12 | 6 | 8 | 26 |
| elongated nectomonad | 14 | 20 | 18 | 18 | 48 | 72 | 12 | 18 | 58 | 44 | 6 |
| short nectomonads | 70 | 6 | 72 | 68 | 34 | 18 | 62 | 66 | 34 | 42 | 66 |
| metacyclic promastigote | 6 | 0 | 4 | 10 | 8 | 8 | 12 | 4 | 2 | 6 | 2 |
